# Cytoplasmic sharing through apical membrane remodeling

**DOI:** 10.1101/2020.02.22.960187

**Authors:** Nora G. Peterson, Benjamin M. Stormo, Kevin P. Schoenfelder, Juliet S. King, Rayson R. S. Lee, Donald T. Fox

## Abstract

Multiple nuclei sharing a common cytoplasm are found in diverse tissues, organisms, and diseases. Yet, multinucleation remains a poorly understood biological property. Cytoplasm sharing invariably involves plasma membrane breaches. In contrast, we discovered cytoplasm sharing without membrane breaching in highly resorptive *Drosophila* rectal papillae. During a six-hour developmental window, 100 individual papillar cells assemble a multinucleate cytoplasm, allowing passage of proteins of at least 27kDa throughout papillar tissue. Papillar cytoplasm sharing does not employ canonical mechanisms such as failed cytokinesis or muscle fusion pore regulators. Instead, sharing requires gap junction proteins (normally associated with transport of molecules <1kDa), which are positioned by membrane remodeling GTPases. Our work reveals a new role for apical membrane remodeling in converting a multicellular epithelium into a giant multinucleate cytoplasm.

**ONE SENTENCE SUMMARY:** Apical membrane remodeling in a resorptive *Drosophila* epithelium generates a shared multinuclear cytoplasm.

## MAIN TEXT

Sharing of cytoplasm in a multinucleate tissue or organism is an important and recurring adaptation across evolution. Multinucleate structures include animal skeletal muscle, mammalian osteoclasts, and mammalian syncytial placental trophoblasts (*1–3*). In disease, cytoplasm sharing facilitates the spread of pathogens (*4*), oncogenic factors (*5, 6*), and prion-like proteins (*7*). Cytoplasm sharing can occur through cytokinesis failure, or through plasma membrane breaches such as fusion pores, tunneling nanotubes, or plasmodesmata. Such clearly visible breaches enable exchange of cytoplasmic components such as RNA, proteins, and even organelles (*8, 9*). The ubiquity and importance of cytoplasm sharing led us to seek out novel examples in the tractable animal model *Drosophila melanogaster.* Here, we report an animal-wide screen for tissues that share cytoplasm. We identify a novel mechanism of cytoplasm sharing in the rectal papilla, a resorptive intestinal epithelium (*10*) and known site of pathogen localization (*11*). Unlike all known examples of multinucleation, cytoplasm sharing in rectal papillae involves developmentally programmed apical membrane remodeling.

To identify new examples of adult tissues in *Drosophila* that share cytoplasm, we ubiquitously expressed *UAS-dBrainbow* (*12*) (**Fig1A**), a Cre-Lox-based system that randomly labels cells with only one of three fluorescent proteins. Multi-labeled cells should only arise by fusion of cells not related by cell division/cytokinesis failure (**Fig1B**). We examined a wide range of tissues (**FigS1A**). From our screen, we discovered that the rectal papilla is a new example of a tissue with cytoplasm sharing. Adult *Drosophila* contain four papillae, each with 100 nuclei, that reside in the posterior hindgut (**Fig1C**). Using both fixed and live imaging of whole organs, we found that at 62 hours post puparium formation (HPPF), each papillar cell contains only one dBrainbow label (**Fig1D**). By contrast, at 69HPPF, multi-labeled cells are apparent (**Fig1D’,F-F’**). We quantitatively measured papillar sharing across the tissue (**FigS1B**, *Methods*) and found that cytoplasm sharing initiates over a narrow 6-hour period (68-74HPPF, **Fig1E**). Our results suggested that at least RNA and possibly protein passes between papillar cells to facilitate cytoplasm sharing. To directly test if protein is shared, we photo-activated GFP (PA-GFP) in single adult papillar cells and observed in real time whether GFP spreads to adjacent cells. We find the principal papillar cells, but not the secondary cells at the papillar base ((*13*), **FigS1C**), share protein across an area of at least several nuclei (**Fig1G-H**). Therefore, proteins as large as ∼27kDa (the size of GFP) can move across an area covered by multiple papillar nuclei. These results indicate that papillae undergo a developmentally programmed conversion from 100 individual cells to a single multinuclear cytoplasm.

**Figure 1.**
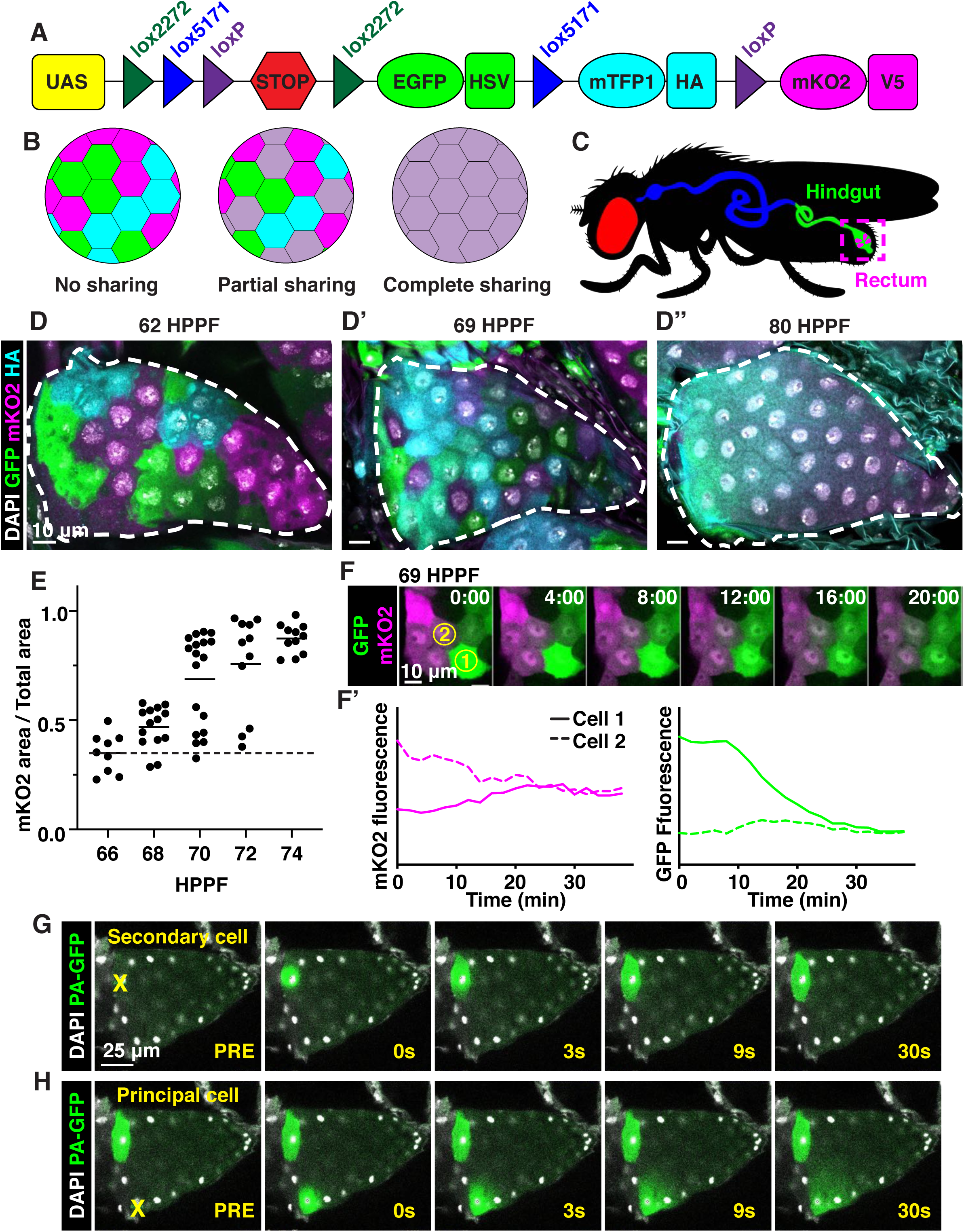
Developmentally programmed cytoplasmic sharing in *Drosophila* papillae. (**A**) dBrainbow. Cre recombinase randomly excises one pair of lox sites, and approximately 1/3 of cells express either EGFP, mKO2, or mTFP1. (**B**) Model of dBrainbow expression with no, partial, or complete cytoplasmic sharing. (**C**) *Drosophila* digestive tract with rectum containing four papillae labeled in magenta box. (**D-D’’**) Representative *dBrainbow* papillae at 62 (**D**), 69 (**D’**), or 80 (**D’’**) hours post-puparium formation (HPPF). (**E**) Cytoplasmic sharing quantification during pupal development. Lines= mean at each time, which differs significantly between 66 and 74 HPPF (p<0.0001). Each point=1 animal (N=9-18, rep=2). (**F**) Live *dBrainbow*-labelled papillar cells during cytoplasmic sharing (69 HPPF). (**F’**) Fluorescence of neighboring cells in (**F**). (**G-H**) Representative adult papilla expressing photo-activatable GFP (PA-GFP). Single cells were photo-activated (yellow X) in secondary cells (**G**) and principal cells (**H**). Time=seconds after activation.

We next examined whether cytoplasm sharing requires the distinctive papillar cell cycle program, which completes prior to sharing onset (**FigS1D**). Larval papillar cells first undergo endocycles, which increase cellular ploidy, and then pupal papillar cells undergo polyploid mitotic cycles, which increase cell number (*14*). Knockdown of the endocycle regulator *fizzy-related* (*15*) significantly disrupts cytoplasm sharing (**FigS1E,F,H**). We hypothesize that endocycles are required for differentiation of the papillae, which later enables these cells to trigger cytoplasm sharing. In contrast, blocking Notch signaling, which initiates papillar mitotic divisions (*14*), does not prevent sharing (**FigS1E,G,H**). Thus, papillar cytoplasm sharing requires developmentally programmed endocycles but not mitotic cycles.

As our *dBrainbow* approach only identifies cytoplasm sharing events that do not involve failed division/cytokinesis, we examined whether sharing results from fusion pore formation, as in skeletal muscle. A well-studied model of such cell-cell fusion in *Drosophila* is myoblast fusion, which requires an actin-based podosome (*16, 17*). We conducted a candidate *dBrainbow*-based RNAi screen (77 genes, **Fig2A, Table S1**) of myoblast fusion regulators and other plasma membrane components. Remarkably, 0/15 myoblast fusion genes from our initial screen regulate papillar cytoplasm sharing (**Fig2A**, **FigS2A**, **Table S1**). Furthermore, dominant-negative forms of Rho family GTPases have no impact on Brainbow labeling (**FigS2B**), providing additional evidence against actin-based cytoplasm sharing. Instead, we found 8/77 genes, including subunits of the vacuolar H+ ATPase (*Vha16-1*), ESCRT-III complex (*Vps2*), and exocyst (*Exo84*) (**Fig2A**) are required for papillar cytoplasm sharing. Through additional screening, the only myoblast fusion regulator required for papillar cytoplasm sharing is *singles bar* (*sing*), a presumed vesicle trafficking gene (*18*) (**FigS2A**). Given the enrichment of our candidate screen hits in membrane trafficking and not myoblast fusion, we further explored the role of membrane trafficking in cytoplasm sharing.

**Figure 2.**
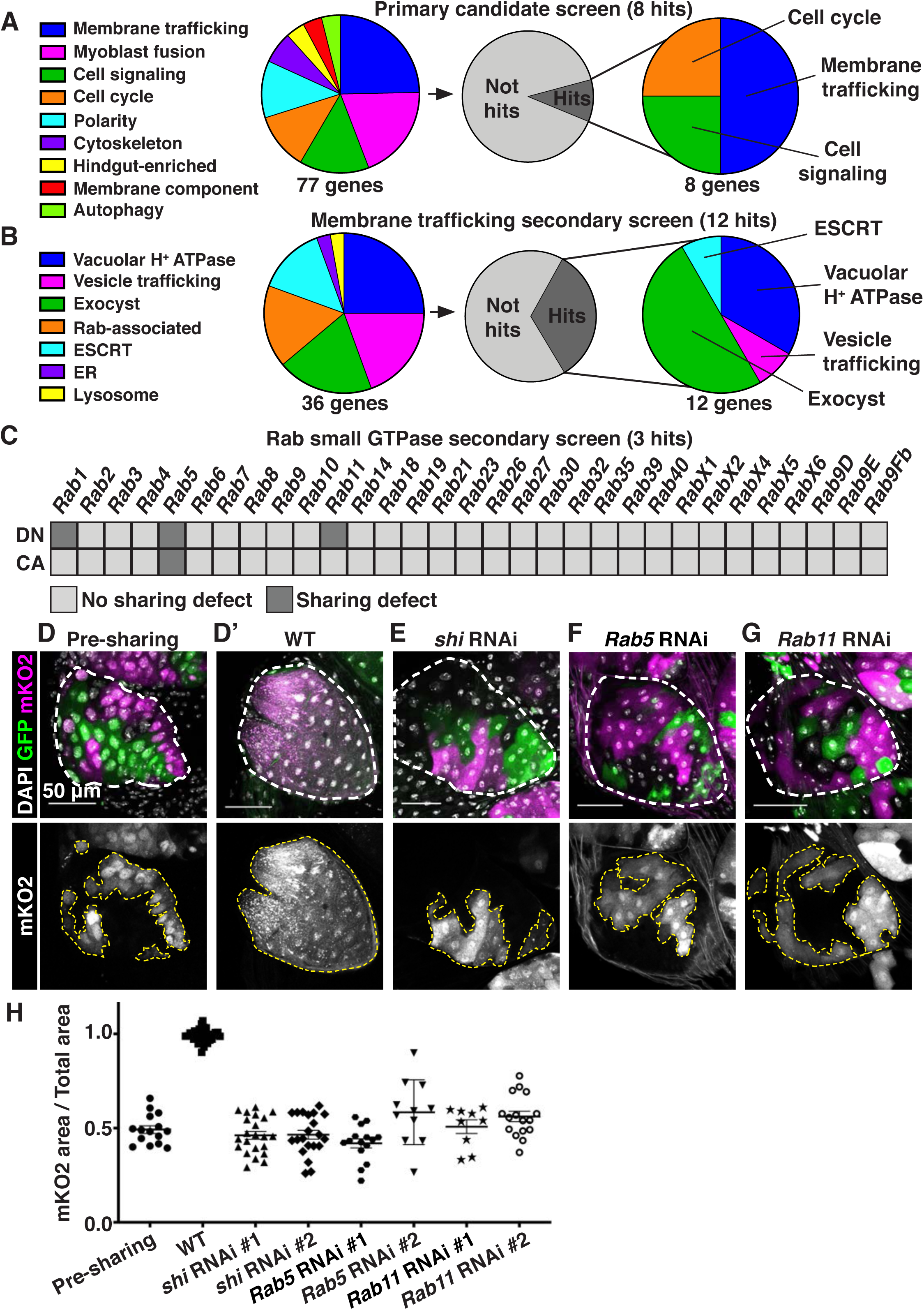
Cytoplasmic sharing requires membrane remodeling proteins. (**A**) Primary *dBrainbow* candidate screen. RNAi and dominant-negative versions of 77 genes representing the indicated roles were screened for sharing defects, and 8 genes were identified. (**B**) Secondary membrane trafficking screen. 36 genes were screened with 12 sharing genes identified. (**C**) Secondary screen of dominant-negative and constitutively-active Rab GTPases. (**D-G**) Representative *dBrainbow* in (**D-D’**) wild type (WT) (**D**) pre-sharing (48HPPF) and (**D’**) post-sharing (young adults), (**E**) adult *shi RNAi,* (**F**) adult *Rab5 RNAi*, (**G**) adult *Rab11 RNAi*. (**H**) Quantification of **D-G**, including two RNAi lines for *shi*, *Rab5*, and *Rab11*. Pre-sharing and knock downs differ significantly from post-sharing WT (p<0.0001, N=9-32, rep=2-3).

We conducted two secondary dBrainbow screens to find specific membrane trafficking pathway components that regulate papillar sharing. First, a focused candidate membrane trafficking screen revealed additional components (12/36 genes screened, **Fig2B, Table S2**) including 3 more vacuolar H+ ATPase subunits, 5 more exocyst components, and the Dynamin GTPase *shibire* (*shi*) (**Fig2B,D,E,H**). Second, we screened constitutively-active and dominant-negative versions of all 31 *Drosophila* Rabs. Sharing requires only a small number of Rabs, specifically the ER/Golgi-associated *Rab1,* the early endosome-associated *Rab5,* and the recycling endosome-associated *Rab11* (**Fig2C,D,F-H**). Given our identification of the membrane vesicle recycling circuit involving *shi*, *Rab5*, and *Rab11*, we focused on these genes. Two unique RNAi lines for each gene show consistent sharing defects, and most of these knockdowns completely recapitulate the pre-sharing state (**Fig2H**). Despite exhibiting strong cytoplasm sharing defects, *shi*, *Rab5*, and *Rab11 RNAi* papillae appear morphologically normal, with only minor cell number decreases (**FigS2C**). These results suggest that membrane recycling GTPases regulate a specific developmental event associated with cytoplasm sharing, and not papillar morphogenesis. In agreement with these GTPases acting during development, rather than as part of an ongoing transport process, GTPase knockdown after sharing onset does not block cytoplasm sharing (**FigS2D-F**). Together, our screens reveal that membrane trafficking, particularly Dynamin-mediated endocytosis and early/recycling endosome trafficking, regulates papillar cytoplasmic sharing.

To better understand how membrane trafficking GTPases initiate cytoplasm sharing during development, we examined endosome and Shi localization during sharing onset. We imaged a GFP-tagged pan-endosome marker (*myc-2x-FYVE*) and a Venus-tagged *shi* before and after sharing. Endosomes are evenly distributed shortly before sharing, but become highly polarized at the basal membrane around the time of sharing onset (**Fig3A-A’,C, FigS3A**). This basal endosome repositioning requires Shi (**Fig3B-C, FigS3A**) and the change in endosome localization is attributed to Rab5-positive early endosomes (**FigS3B-C**). Additionally, Shi localization changes from apical polarization to a uniform distribution during sharing onset (**Fig3D-E**). These localization changes indicate that membrane trafficking factors are dynamic during cytoplasm sharing onset.

**Figure 3.**
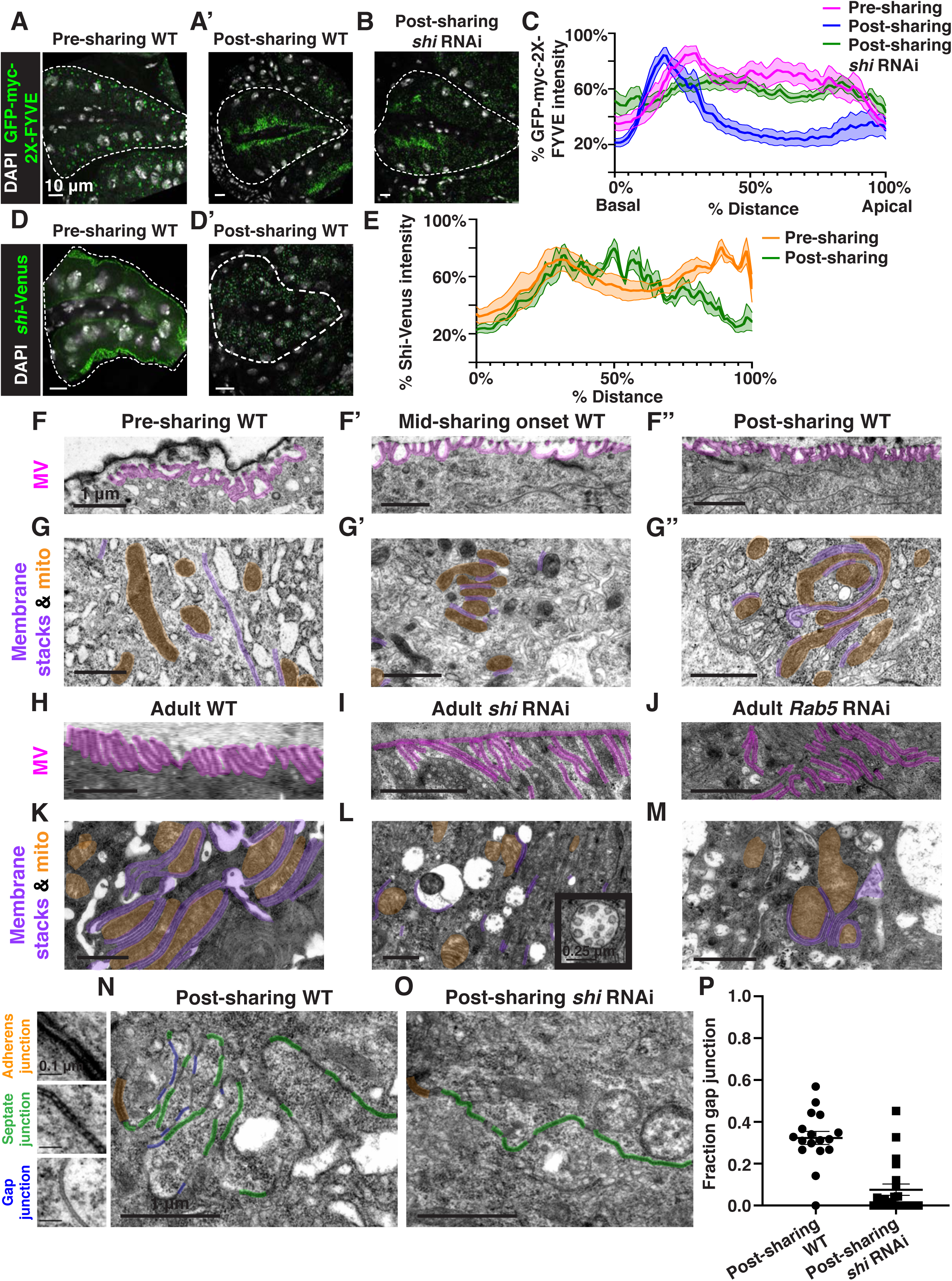
Gap junction establishment, but no membrane breaches, accompany cytoplasm sharing. (**A-A’**) Endosome localization (GFP-myc-2x-FYVE), representative of (**A**) pre- and (**A’**) post-sharing onset. (**B**) Endosomes in *shi RNAi* post-sharing, *see Methods*. (**C**) Aggregated endosome line profiles for WT pre-sharing (N=6, rep=3), WT post-sharing (N=7, rep=2), and *shi RNAi* post-sharing (N=10, rep=2). Shaded area represents standard error. (**D-D’**) Shi-Venus localization pre- and post-sharing onset. (**E**) Line profiles as in (**D-D’**) (N=4-5, rep=3). (**F-O**) Representative Transmission Electron Micrographs (TEMs). **(F-F’’**) Microvillar-like structures (MV) pre (**F**), mid-(**F’**), and post-(**F’’**) sharing onset. (**G-G’’**) Mitochondria and surrounding membrane pre-(**G**), mid-(**G’**), and post-(**G’’**) sharing onset. (**H-J**) Microvillar-like structures (MV) of adult papillae in WT (**H**), *shi RNAi* (**I**), and *Rab5 RNAi* (**J**). (**K-M**) Mitochondria and surrounding membranes of adult papillae in WT (**K**), *shi RNAi* (**L**), and *Rab5 RNAi* (**M**). Inset in **L** shows trapped vesicles. (**N-O**) WT and *shi RNAi* post-sharing. Adherens (orange), septate (green), and gap (blue) junctions are highlighted. (**P**) Quantification of the ratio of gap junction length to septate plus gap junction length (Fraction gap junction) (N=3-4, rep=2). p<0.0001 for the difference in gap junction ratio between WT and *shi RNAi*.

To determine what membrane remodeling events underlie GTPase-dependent cytoplasm sharing, we turned to ultrastructural analysis. Adult ultrastructure and physiology of papillar cells has been examined previously in *Drosophila* (*19*) and related insects (*20*). These cells contain elaborate membrane networks that facilitate selective ion resorption from the gut lumen, facing the apical side of papillar cells, to the hemolymph, facing the basal side. Still, little is known about developmental processes or mechanisms governing the unique papillar cell architecture. We looked for changes in cell-cell junctions and lateral membranes that coincide with cytoplasm sharing, especially to determine if there is a physical membrane breach between cells. We identified several dramatic changes in membrane architecture. First, apical microvilli-like structures form during sharing onset (**Fig3F-F’’**). Just basal to the microvilli, apical cell-cell junctions compress from a straight to a more tortuous morphology around the time of cytoplasm sharing onset **(FigS3D-D’’**). One of the most striking changes, coincident with Shi re-localization, is formation of pan-cellular endomembrane stacks surrounding mitochondria. These stacks are likely ion transport sites (**Fig3G-G’’**). Thus, massive apical and intracellular plasma membrane reorganization coincides with both cytoplasm sharing and Shi/endosome re-localization. We next assessed whether the extensive membrane remodeling requires Shi, Rab5, and Rab11. In *shi* and *Rab5 RNAi* animals, microvilli protrude downward, instead of upward (**Fig3H-J**). Additionally, apical junctions do not compress as in controls (**FigS3E-G**). Notably, membrane stacks are greatly reduced (**Fig3K-M**). *shi RNAi* animals exhibit numerous trapped vesicles, consistent with a known role for Dynamin in membrane vesicle severing (*21, 22*) (**Fig3L, inset**). Together, we find that Shi and endosomes extensively remodel membranes during cytoplasm sharing.

Our extensive ultrastructural analysis did not reveal any clear breaches in the plasma membrane, despite numerous membrane alterations. Adult papillae exhibit large extracellular spaces between nuclei that eliminate the possibility of cytoplasm sharing throughout much of the lateral membrane (**FigS4A**) (*19, 20*). Instead, through our GTPase knockdown studies, we identified a striking alteration in the apical cell-cell interface that strongly correlates with cytoplasm sharing. Specifically, *shi* animals frequently lack apical gap junctions (**Fig3N-O**) (p<0.0001) (**Fig3P**, **FigS3H-H’’**). Upon closer examination of control animal development, we find that apical gap junction-like structures arise at cytoplasm sharing onset. There is almost no gap junction-like structure before cytoplasm sharing (**Fig4A-B**, **FigS5A-A’’**). Given our electron micrograph results, we determined which innexins, the protein family associated with gap junctions in invertebrates (*23, 24*), are expressed in rectal papillae. From RNA-seq data (*Methods*), we determined that *ogre* (*Inx1*), *Inx2*, and *Inx3* are most highly expressed (**Fig4C**). This combination of innexins is not unique; the non-sharing brain and optic lobe (**FigS1A**) also express high levels of all three (*25*). We examined localization of Inx3 (a gap junction component), and compared it to a septate junction component, NeurexinIV (NrxIV). NrxIV localizes similarly both pre and post-sharing onset (**Fig4D-D’**), indicative of persistent septate junctions remaining between papillar cells. In contrast, Inx3 organizes apically only after cytoplasm sharing (**Fig4E-E’**, **FigS5B-B’**). We tested whether innexins are required for cytoplasm sharing. Knocking down these three genes individually causes mild yet significant cytoplasm sharing defects (**Fig4F**). However, we see larger defects when we express dominant-negative *ogre^DN^* (**Fig4F-G**), which contains a C-terminal GFP tag that interferes with channel passage. Also, heterozygous animals containing a ten gene-deficiency spanning *ogre*, *Inx2*, and *Inx7* have more severe defects (**Fig4F**, *Df(1)BSC867*). Finally, we tested whether cytoplasm sharing is essential for normal rectal papillar function. Rectal papillae selectively absorb water and ions from the gut lumen for transport back into the hemolymph, and excrete unwanted lumen contents. One test of papillar function is viability following the challenge of a high-salt diet (*15, 26*). Using either pan-hindgut or papillae-specific (**FigS5C-D,** *Methods*) knockdown of cytoplasm sharing regulators, we find both *shi^DN^* and *ogre^DN^* animals are extremely sensitive to the high-salt diet (mean survival <1 day, **Fig4H**). These results underscore an important function for gap junction proteins, as well as membrane remodeling by Shibire, in cytoplasm sharing.

**Figure 4.**
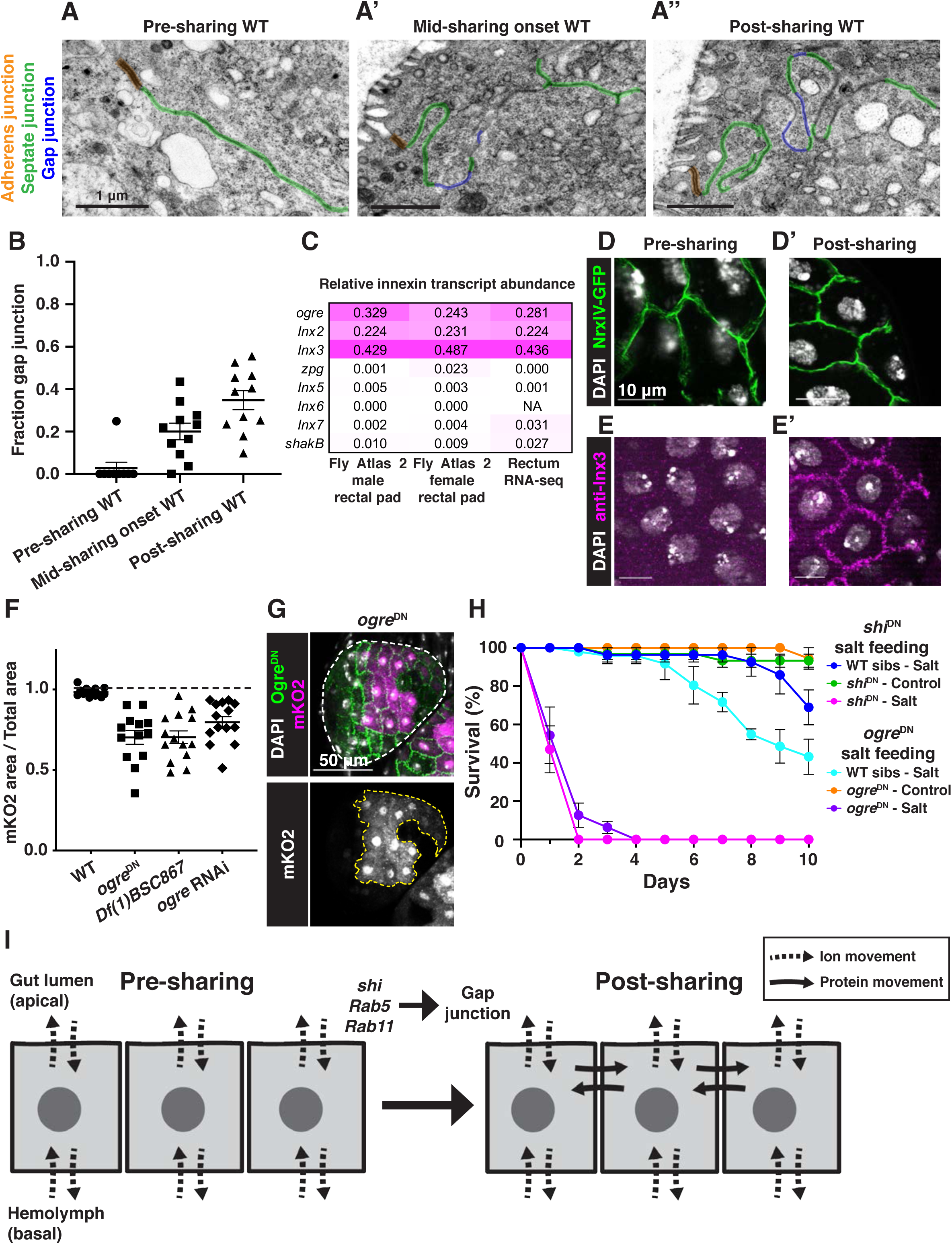
Gap junction proteins are required for cytoplasmic sharing. (**A-A’’**) Representative apical junctions highlighted by junctional type in pre (**A**), mid (**A’**), and post (**A’’**) sharing onset. (**B**) Quantification of fraction gap junction (gap junction length / (gap + septate junction length)) in pre-, mid-, and post-sharing onset pupae (N=3-4, rep=2). (**C**) *Drosophila* innexin expression in the adult rectum (*Methods*). (**D-D’**) Adherens junctions in pre-(**D**) and post-(**D’**) sharing pupae visualized by *NrxIV-GFP*. (E-E’) WT pupae pre- and post-sharing onset stained with anti-Inx3. (**F**) Quantification of cytoplasm sharing in WT, *ogre^DN^*, *Df(1)BSC867/+* (a 10-gene deficiency covering *ogre*, *Inx2*, and *Inx7*), and *ogre RNAi* adult papillae (N=13-14, rep=2). (**G**) Representative adult rectal papilla expressing *GFP-ogre* and *dBrainbow*. (**H**) Survival of WT, *shi^DN^*, and *ogre^DN^* animals on a high-salt diet (N=27-37, rep=3). (**I**) Proposed model for cytoplasmic sharing in an intact papillar epithelium.

Our findings identify *Drosophila* rectal papillae as a new and distinctive example of cytoplasm sharing in a simple, genetically tractable system. Papillar cytoplasm sharing is developmentally regulated, occurring over a brief 6-hour window, and requires membrane remodeling by trafficking GTPases, which apically position gap junction proteins (**Fig4I, FigS5H**). These membrane and junctional changes are required for normal rectum function. We speculate that papillar cytoplasm movement across a giant multinuclear structure enhances resorption by facilitating interaction of ions and ion transport machinery with intracellular membrane stacks. Given the absence of other clear canals, channels, or breaks in lateral membrane, our data suggest a specialized function of gap junction proteins facilitates cytoplasm sharing between neighboring cells in an otherwise intact epithelium (**Fig4I**). Although gap junctions typically transfer molecules of <1kDa, elongated proteins up to 18 kDa are observed to pass through certain vertebrate gap junctions (*27*). Our results have several implications for functions and regulation of multinucleation. Given that cytoplasm sharing facilitates pathogen spread (*4*), and that papillae are an avenue of entry for mosquito viruses (*11*), our findings may impact insect vector control strategies. Our prior work (*15*) revealed that papillae are highly tolerant of chromosome mis-segregation, and our work here suggests this tolerance may be due in part to neutralization of aneuploidies through cytoplasm sharing, a finding relevant to syncytial cancers. In the future, our Brainbow-based approach could be applied to other contexts to identify other tissues with gap junction-dependent but membrane breach-independent cytoplasm sharing. Collectively, our findings highlight the expanding diversity of multicellular tissue organization strategies.

## ACKNOWLEDGMENTS

We thank members of the Fox laboratory and Drs. Dong Yan and Tony Harris for valuable feedback. Ying Hao (Duke Eye Center) provided assistance with electron microscopy.

## Author contributions

experimental design, execution, and data analysis-all authors, manuscript writing and editing-N.P. and D.F.

## Data and materials availability

all data is available in the main text or the supplementary materials.

## SUPPLEMENTARY MATERIAL

### MATERIALS AND METHODS

#### Fly Stocks and Genetics

Flies were raised at 25C on standard media (Archon Scientific, Durham, NC) unless specified otherwise. See **Table S4** for a list of fly stocks used. See **Table S3** for a full list of fly lines screened in primary and secondary screens. See **Table S5** for panel-specific genotypes.

*brachyenteron (byn)-Gal4* was the driver for all UAS transgenes with the exception of the screen in FigS1A, which used *Tubulin-Gal4*, and the *shi* knockdown in Fig4H, which used *60H12-Gal4*. *60H12-Gal4* expresses only in the papillar cells and not the rest of the hindgut, and use of this driver blocks cytoplasm sharing using *UAS-shi^DN^* (FigS5C-G). For all *Gal4* experiments, *UAS* expression was at 29C, except in Fig1D-F, where it was at 25C. If *byn-Gal4* expression of a given *UAS-*transgene was lethal, the experiment was repeated with a temperature-sensitive *Gal80^ts^* repressor transgene and animals were kept at 18C until shifting to 29C at an experimentally-determined time point that would both result in viable animals and permit time to express the transgene prior to syncytium formation.

For salt feeding assays, age- and sex-matched siblings were transferred into vials containing 2% NaCl food made with Nutri-Fly MF® food base (Genesee Scientific) or control food (*15*). Flies were monitored for survival each day for 10 days.

#### Tissue Preparation

For fixed imaging, tissues were dissected in PBS and immediately fixed in 3.7% formaldehyde + 0.3% Triton-X for 15 minutes. Immunostaining was performed in 0.3% Triton-X with 1% normal goat serum (*14*). The following antibodies were used: Rabbit anti-GFP (Thermo-Fisher, Waltham MA, A11122, 1:1000), Rat anti-HA (Sigma, 3F10, 1:100), Rabbit anti-Inx3 (generous gift from Reinhard Bauer, 1:75, (*27*)), 488, 568, 633 secondary antibodies (Life Technologies, Alexa Fluor ®, 1:2000). Tissue was stained with DAPI at 5μg/ml and mounted in VECTASHIELD Mounting Media on slides.

#### Microscopy

##### Light Microscopy

For fixed imaging, images were obtained on either a Leica SP5 inverted confocal with a 40X/1.25NA oil objective with emission from a 405 nm diode laser, a 488 nm argon laser, a 561 nm Diode laser, and a 633 HeNe laser under control of Leica LAS AF 2.6 software, or on an Andor Dragonfly Spinning Disk Confocal plus. Images were taken with two different cameras, iXon Life 888 1024 x 1024 EMCCD (pixel size 13um) and the Andor Zyla PLUS 4.2 Megapixel sCMOS 2048 x 2048 (pixel size 6.5um) depending on imaging needs. Images were taken on the **40x**/1.25-0.75 oil 11506250: 40X, HCX PL APO, NA: 1.25, Oil, DIC, WD: 0.1mm, coverglass: 0.17mm, Iris diaphragm, Thread type: M25, **63x**/1.20 water 11506279: 63X, HCX PL APO W Corr CS, NA: 1.2, Water, DIC, WD: 0.22mm, Coverglass: 0.14mm-0.18mm, thread type: M25, and **100x**/1.4-0.70 oil 11506210: HCX PL APO, NA: 1.4, Oil, DIC, WD: 0.09mm, Coverglass: 0.17mm, Iris Diaphragm, Thread type: M25. The lasers used were: 405nm diode laser, 488nm argon laser, 561nm diode laser, and HeNe 633nm laser.

For live imaging, hindguts were dissected and cultured based on previous protocols (14). Live imaging of cell fusion was performed on a spinning disc confocal (Yokogawa CSU10 scanhead) on an Olympus IX-70 inverted microscope using a 40x/1.3 NA UPlanFl N Oil objective, a 488 nm and 568 nm Kr-Ar laser lines for excitation and an Andor Ixon3 897 512 EMCCD camera. The system was controlled by MetaMorph 7.7.

Photo-activation was carried out using Leica SP5 and SP8 microscopes and the FRAP Wizard embedded in the Leica AS-F program. An initial z-stack of the tissue was acquired both before and after activation to examine the full extent of PA-GFP movement in three dimensions. PA-GFP was activated by either point activation or region of interest activation with the 405nm laser set to between 5-20%, depending on the microscope and sample of interest. For each imaging session, test activations on nearby tissues were performed prior to quantified experiments to ensure that only single cells were being activated. After activation, the wizard software was used to acquire time lapses of 15s-2min of a single activation plane in order to capture protein movement. Extremely low 488nm and 405nm laser power was used in acquisition of the time lapse images of GFP and Hoechst respectively. Low level 405nm scanning did not significantly activate PA-GFP, and control experiments were performed without the use of 405nm time lapses and showed the same protein movement results (data not shown).

##### Transmission Electron Microscopy

Hindguts were dissected into PBS and fixed in a solution of 2.5% glutaraldehyde in 0.1% cacodylate buffer, pH 7.2. Post-fix specimens were stained with 1% osmium tetroxide in 0.1M cacodylate buffer, dehydrated, soaked in a 1:1 propylene oxide:Epon 812 resin, and then embedded in molds with fresh Epon 812 resin at 65C overnight. The blocks were cut into semi-thin (0.5µm) sections using Leica Reichert Ultracuts and the sections were stained with 1% methylene blue. After inspection, ultra-thin sections (65nm-75nm) were cut using Leica EM CU7 and contrast stained with 2% uranyl acetate, 3.5% lead citrate solution. Ultrathin sections were visualized on a JEM-1400 transmission electron microscope (JEOL) using an ORIUS (1000) CCD 35mm port camera.

#### Image Analysis

All image analysis was performed using ImageJ and FIJI (*28, 29*).

##### Cytoplasm sharing calculation

Cytoplasmic sharing was quantified by manually tracing the total papillar area by morphology and the area marked by mKO2 signal in one z-slice of the papillar face of each animal. The area marked by mKO2 was summed and divided by the sum of the total papillar area to yield the papillar fraction marked by mKO2 which indicates the degree of cytoplasmic sharing within each animal. Papillae without mKO2 signal were excluded from the area measurements.

##### Line profiles

For line profile data collection, fixed and mounted hindguts were imaged on a Zeiss Apotome on the 40Xoil objective. Once moved into ImageJ, the images were rotated with no interpolation so that the central canal was perpendicular to the bottom of the image. From the midline of the central canal, a straight line (width of 300) was drawn out to one edge of the papillae. One papilla was measured per animal. Papillae were measured at the widest width. Next, the Analyze > Plot Profile data was collected from this representative 300 width line and moved into Excel. In Excel, the data was first was normalized to the maximum length of the papillae and the maximum GFP intensity per animal. Each data point is a % of the total length of the papillae and a % of the maximum GFP intensity. Next, the X values were rounded to its nearest 1% value. Next, all the Y-values were averaged per X value bins (average % GFP intensity per rounded % distance value). % GFP intensity values were plotted from 1-100% total distance of papilla.

## Genotype and experiment-specific method notes

Some additional methodological details, including animal genotype, applied to only a specific figure panel. Please see **Table S5** for this information.

**Table S1.** Cytoplasm sharing primary candidate screen gene results.

| Gene category | Gene | Annotation symbol | Gene ID | Sharing disrupted? |
| --- | --- | --- | --- | --- |
| Autophagy | *Atg1* | CG10967 | FBgn0260945 | No |
| Autophagy | *Atg7* | CG5489 | FBgn0034366 | No |
| Autophagy | *Atg8a* | CG32672 | FBgn0052672 | No |
| Cell cycle / Chromosomes | *blue* | NA | FBgn0283709 | No |
| Cell cycle / Chromosomes | *CapD2* | CG1911 | FBgn0039680 | No |
| Cell cycle / Chromosomes | *Cdc2* | CG5363 | FBgn0004106 | Yes |
| Cell cycle / Chromosomes | *Clamp* | CG1832 | FBgn0032979 | No |
| Cell cycle / Chromosomes | *endos* | CG6513 | FBgn0061515 | No |
| Cell cycle / Chromosomes | *fzr* | CG3000 | FBgn0262699 | Yes |
| Cell cycle / Chromosomes | *Mi-2* | CG8103 | FBgn0262519 | No |
| Cell cycle / Chromosomes | *Rbp9* | CG3151 | FBgn0010263 | No |
| Cell cycle / Chromosomes | *SA-2* | CG13916 | FBgn0043865 | No |
| Cell signaling | *Chico* | CG5686 | FBgn0024248 | No |
| Cell signaling | *Egfr* | CG10079 | FBgn0003731 | Yes |
| Cell signaling | *grk* | CG17610 | FBgn0001137 | No |
| Cell signaling | *N* | CG3936 | FBgn0004647 | No |
| Cell signaling | *Ptp61F* | CG9181 | FBgn0267487 | No |
| Cell signaling | *rho* | CG1004 | FBgn0004635 | Yes |
| Cell signaling | *ru* | CG1214 | FBgn0003295 | No |
| Cell signaling | *spi* | CG10334 | FBgn0005672 | No |
| Cell signaling | *stet* | CG33166 | FBgn0020248 | No |
| Cell signaling | *wts* | CG12072 | FBgn0011739 | No |
| Cell signaling | *βggt-II* | CG18627 | FBgn0028970 | No |
| Cytoskeleton | *ALiX* | CG12876 | FBgn0086346 | No |
| Cytoskeleton | *Cdc42* | CG12530 | FBgn0010341 | No |
| Cytoskeleton | *DCTN1-p150* | CG9206 | FBgn0001108 | No |
| Cytoskeleton | *pav* | CG1258 | FBgn0011692 | No |
| Cytoskeleton | *wash* | CG13176 | FBgn0033692 | No |
| Hindgut-enriched | *dac* | CG4952 | FBgn0005677 | No |
| Hindgut-enriched | *Dr* | CG1897 | FBgn0000492 | No |
| Hindgut-enriched | *nrv3* | CG8663 | FBgn0032946 | No |
| Membrane component | *Flo1* | CG8200 | FBgn0024754 | No |
| Membrane component | *Flo2* | CG32593 | FBgn0264078 | No |
| Membrane component | *Iris* | CG4715 | FBgn0031305 | No |
| Myoblast fusion | *Arf51F* | CG8156 | FBgn0013750 | No |
| Myoblast fusion | *Arp2* | CG9901 | FBgn0011742 | No |
| Myoblast fusion | *Arp3* | CG7558 | FBgn0262716 | No |
| Myoblast fusion | *Ced-12* | CG5336 | FBgn0032409 | No |
| Myoblast fusion | *dock* | CG3727 | FBgn0010583 | No |
| Myoblast fusion | *hbs* | CG7449 | FBgn0029082 | No |
| Myoblast fusion | *Hem* | CG5837 | FBgn0011771 | No |
| Myoblast fusion | *mbc* | CG10379 | FBgn0015513 | No |
| Myoblast fusion | *Rac1* | CG2248 | FBgn0010333 | No |
| Myoblast fusion | *Rho1* | CG8416 | FBgn0014020 | No |
| Myoblast fusion | *rols* | CG32096 | FBgn0041096 | No |
| Myoblast fusion | *rst* | CG4125 | FBgn0003285 | No |
| Myoblast fusion | *SCAR* | CG4636 | FBgn0041781 | No |
| Myoblast fusion | *siz* | CG32434 | FBgn0026179 | No |
| Myoblast fusion | *WASp* | CG1520 | FBgn0024273 | No |
| Polarity | *Abi* | CG9749 | FBgn0020510 | No |
| Polarity | *CadN* | CG7100 | FBgn0015609 | No |
| Polarity | *cindr* | CG31012 | FBgn0027598 | No |
| Polarity | *cno* | CG42312 | FBgn0259212 | No |
| Polarity | *Gli* | CG3903 | FBgn0001987 | No |
| Polarity | *l(2)gl* | CG2671 | FBgn0002121 | No |
| Polarity | *Nrg* | CG1634 | FBgn0264975 | No |
| Polarity | *sdt* | CG32717 | FBgn0261873 | No |
| Polarity | *shg* | CG3722 | FBgn0003391 | No |
| Vesicle trafficking | *Atl* | CG6668 | FBgn0039213 | No |
| Vesicle trafficking | *Bet1* | CG14084 | FBgn0260857 | No |
| Vesicle trafficking | *Chmp1* | CG4108 | FBgn0036805 | No |
| Vesicle trafficking | *CHMP2B* | CG4618 | FBgn0035589 | No |
| Vesicle trafficking | *dnd* | CG6560 | FBgn0038916 | No |
| Vesicle trafficking | *Exo84* | CG6095 | FBgn0266668 | Yes |
| Vesicle trafficking | *lerp* | CG31072 | FBgn0051072 | No |
| Vesicle trafficking | *Rab11* | CG5771 | FBgn0015790 | Yes |
| Vesicle trafficking | *Rab23* | CG2108 | FBgn0037364 | No |
| Vesicle trafficking | *Rab4* | CG4921 | FBgn0016701 | No |
| Vesicle trafficking | *Rab7* | CG5915 | FBgn0015795 | No |
| Vesicle trafficking | *Rab8* | CG8287 | FBgn0262518 | No |
| Vesicle trafficking | *RabX4* | CG31118 | FBgn0051118 | No |
| Vesicle trafficking | *Vha16-1* | CG3161 | FBgn0262736 | Yes |
| Vesicle trafficking | *Vha55* | CG17369 | FBgn0005671 | No |
| Vesicle trafficking | *VhaAC39-1* | CG2934 | FBgn0285910 | No |
| Vesicle trafficking | *VhaAC39-2* | CG4624 | FBgn0039058 | No |
| Vesicle trafficking | *Vps2* | CG14542 | FBgn0039402 | Yes |
| Vesicle trafficking | *Vps33b* | CG5127 | FBgn0039335 | No |
| <u>Total screen results</u> |  |  |  |  |
| <u>Sharing disrupted</u> | 8 |  |  |  |
| <u>No sharing phenotype</u> | 69 |  |  |  |
| **Total** | 77 |  |  |  |
| <u>Screen results by category</u> |  |  |  |  |
| Polarity | 9 |  |  |  |
| Vesicle trafficking | 19 |  |  |  |
| Myoblast fusion | 15 |  |  |  |
| Cell cycle / Chromosomes | 9 |  |  |  |
| Cell signaling | 11 |  |  |  |
| Autophagy | 3 |  |  |  |
| Cytoskeleton | 5 |  |  |  |
| Hindgut-enriched | 3 |  |  |  |
| Membrane component | 3 |  |  |  |
| **Total** | **77** |  |  |  |

**Table S2.** Membrane trafficking primary and secondary candidate screen gene results.

| **Gene category** | **Gene subcategory** | **Gene** | **Annotation symbol** | **Gene ID** | **Sharing disrupted?** | **Screen** |
| --- | --- | --- | --- | --- | --- | --- |
| Membrane trafficking | ER | *Atl* | CG6668 | FBgn0039213 | No | Primary |
| Membrane trafficking | ESCRT | *Chmp1* | CG4108 | FBgn0036805 | No | Primary |
| Membrane trafficking | ESCRT | *CHMP2B* | CG4618 | FBgn0035589 | No | Primary |
| Membrane trafficking | ESCRT | *Isn* | CG6637 | FBgn0260940 | No | Secondary |
| Membrane trafficking | ESCRT | *Vps2* | CG14542 | FBgn0039402 | Yes | Primary |
| Membrane trafficking | ESCRT | *Vps4* | CG6842 | FBgn0283469 | No | Secondary |
| Membrane trafficking | Exocyst | *Exo70* | CG7127 | FBgn0266667 | No | Secondary |
| Membrane trafficking | Exocyst | *Exo84* | CG6095 | FBgn0266668 | Yes | Primary |
| Membrane trafficking | Exocyst | *Sec10* | CG6159 | FBgn0266673 | Yes | Secondary |
| Membrane trafficking | Exocyst | *Sec15* | CG7034 | FBgn0266674 | Yes | Secondary |
| Membrane trafficking | Exocyst | *Sec5* | CG8843 | FBgn0266670 | Yes | Secondary |
| Membrane trafficking | Exocyst | *Sec6* | CG5341 | FBgn0266671 | Yes | Secondary |
| Membrane trafficking | Exocyst | *Sec8* | CG2095 | FBgn0266672 | Yes | Secondary |
| Membrane trafficking | Lysosome | *lerp* | CG31072 | FBgn0051072 | No | Primary |
| Membrane trafficking | Rab-associated | *CG41099* | CG41099 | FBgn0039955 | No | Secondary |
| Membrane trafficking | Rab-associated | *mtm* | CG9115 | FBgn0025742 | No | Secondary |
| Membrane trafficking | Rab-associated | *nuf* | CG33991 | FBgn0013718 | No | Secondary |
| Membrane trafficking | Rab-associated | *Rala* | CG2849 | FBgn0015286 | No | Secondary |
| Membrane trafficking | Rab-associated | *Rep* | CG8432 | FBgn0026378 | No | Secondary |
| Membrane trafficking | Rab-associated | *Rip11* | CG6606 | FBgn0027335 | No | Secondary |
| Membrane trafficking | Vacuolar H<sup>+</sup> ATPase | *Vha16-1* | CG3161 | FBgn0262736 | Yes | Primary |
| Membrane trafficking | Vacuolar H<sup>+</sup> ATPase | *Vha16-2* | CG32089 | FBgn0028668 | No | Secondary |
| Membrane trafficking | Vacuolar H<sup>+</sup> ATPase | *Vha16-3* | CG32090 | FBgn0028667 | No | Secondary |
| Membrane trafficking | Vacuolar H<sup>+</sup> ATPase | *Vha16-5* | CG6737 | FBgn0032294 | Yes | Secondary |
| Membrane trafficking | Vacuolar H<sup>+</sup> ATPase | *Vha55* | CG17369 | FBgn0005671 | No | Primary |
| Membrane trafficking | Vacuolar H<sup>+</sup> ATPase | *VhaAC39-1* | CG2934 | FBgn0285910 | No | Primary |
| Membrane trafficking | Vacuolar H<sup>+</sup> ATPase | *VhaAC39-2* | CG4624 | FBgn0039058 | No | Primary |
| Membrane trafficking | Vacuolar H<sup>+</sup> ATPase | *VhaPPA1-1* | CG7007 | FBgn0028662 | Yes | Secondary |
| Membrane trafficking | Vacuolar H<sup>+</sup> ATPase | *VhaPPA1-2* | CG7026 | FBgn0262514 | Yes | Secondary |
| Membrane trafficking | Vesicle trafficking | *Bet1* | CG14084 | FBgn0260857 | No | Primary |
| Membrane trafficking | Vesicle trafficking | *Chc* | CG9012 | FBgn0000319 | No | Secondary |
| Membrane trafficking | Vesicle trafficking | *dnd* | CG6560 | FBgn0038916 | No | Primary |
| Membrane trafficking | Vesicle trafficking | *shi* | CG18102 | FBgn0003392 | Yes | Secondary |
| Membrane trafficking | Vesicle trafficking | *Vps29* | CG4764 | FBgn0031310 | No | Secondary |
| Membrane trafficking | Vesicle trafficking | *Vps33b* | CG5127 | FBgn0039335 | No | Primary |
| Membrane trafficking | Vesicle trafficking | *Vps35* | CG5625 | FBgn0034708 | No | Secondary |
| <u>Total screen results</u> |  |  |  |  |  |  |
| <u>Sharing disrupted</u> | 12 |  |  |  |  |  |
| <u>No sharing phenotype</u> | 24 |  |  |  |  |  |
| **Total** | 36 |  |  |  |  |  |
| <u>Screen results by category</u> | **Total** | **Hits** |  |  |  |  |
| ER | 1 | 0 |  |  |  |  |
| ESCRT | 5 | 1 |  |  |  |  |
| Exocyst | 7 | 6 |  |  |  |  |
| Lysosome | 1 | 0 |  |  |  |  |
| Rab-associated | 6 | 0 |  |  |  |  |
| Vacuolar H<sup>+</sup> ATPase | 9 | 4 |  |  |  |  |
| Vesicle trafficking | 7 | 1 |  |  |  |  |
| **Total** | 36 |  |  |  |  |  |

**Table S3.**
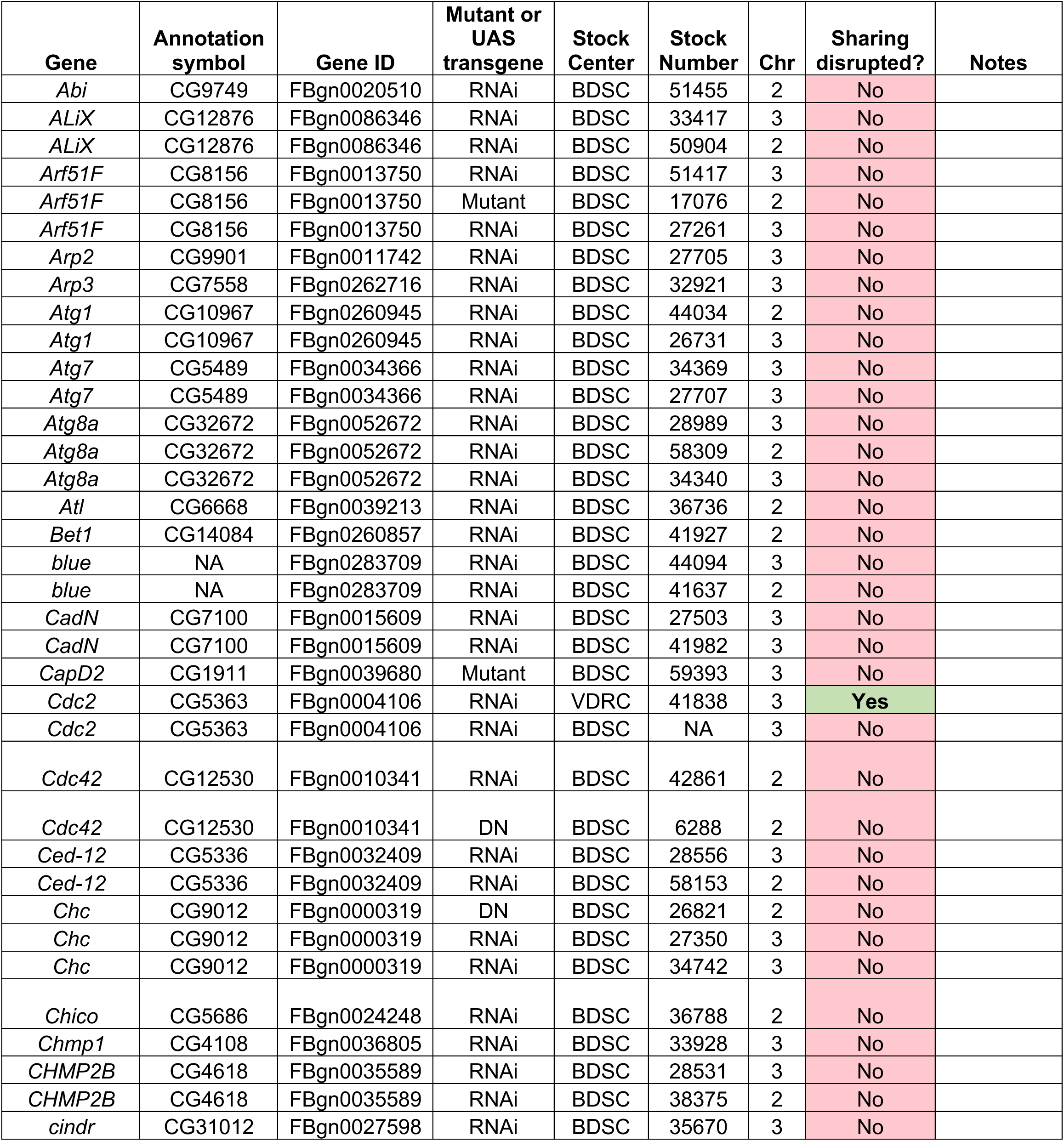
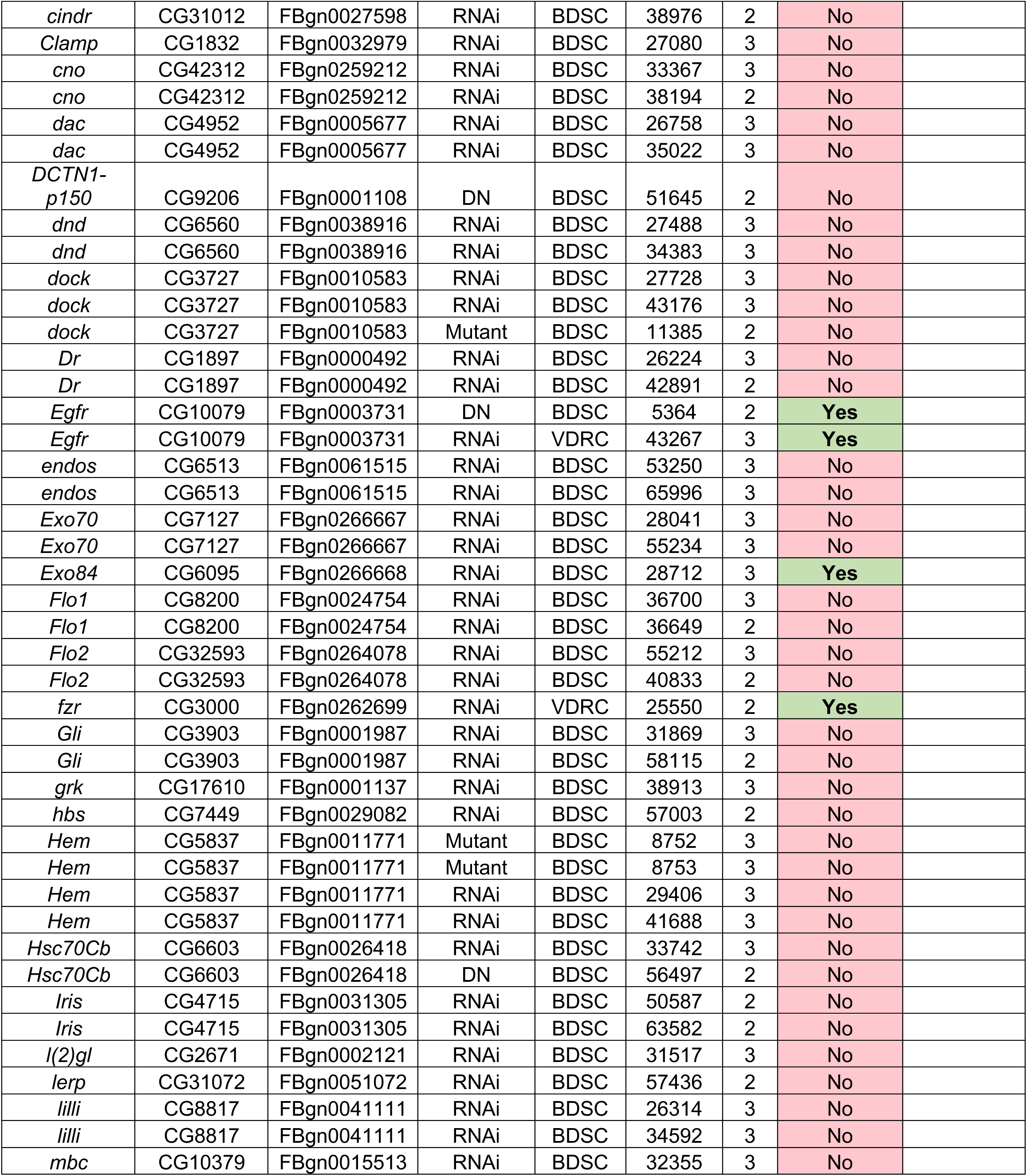
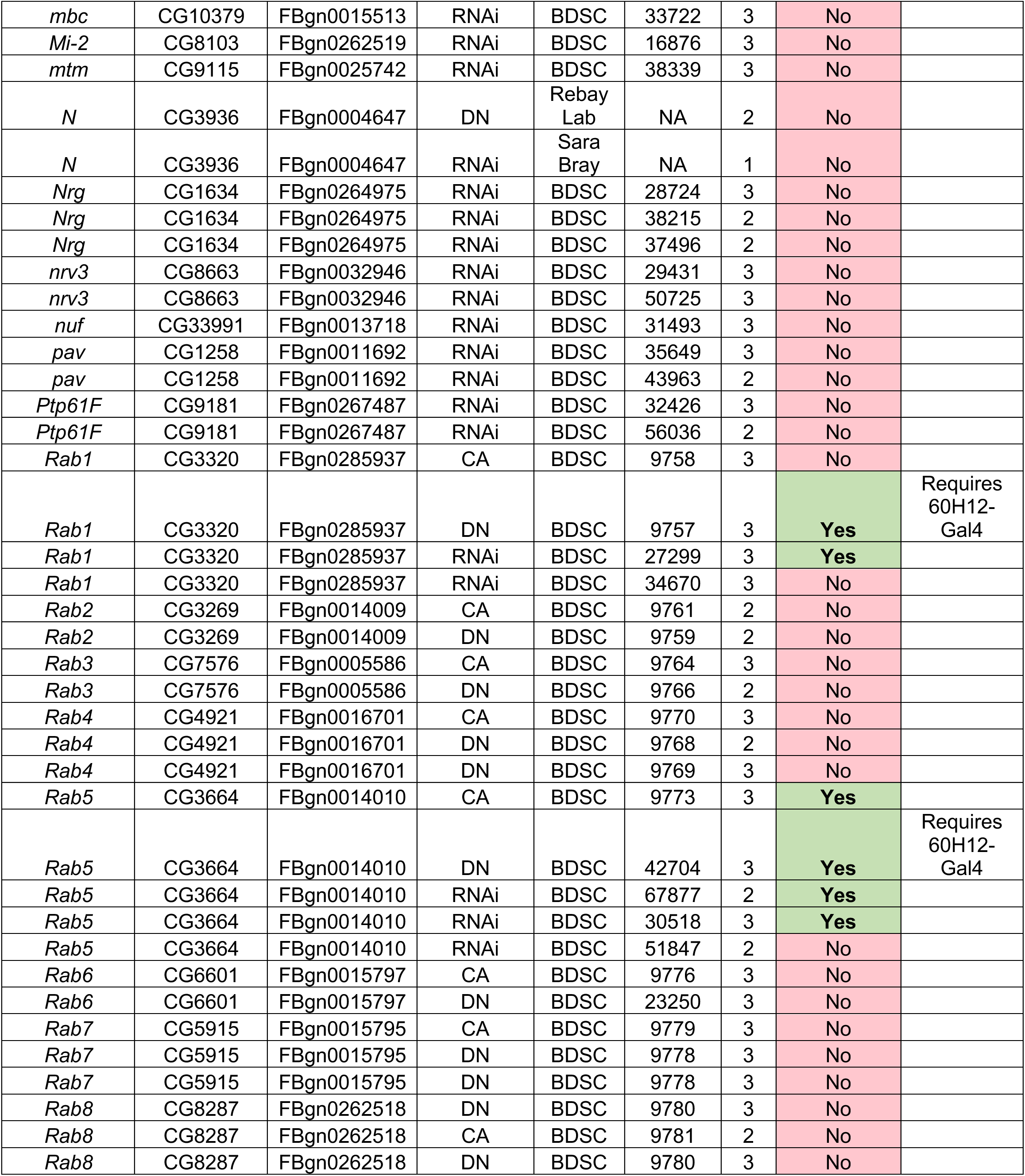
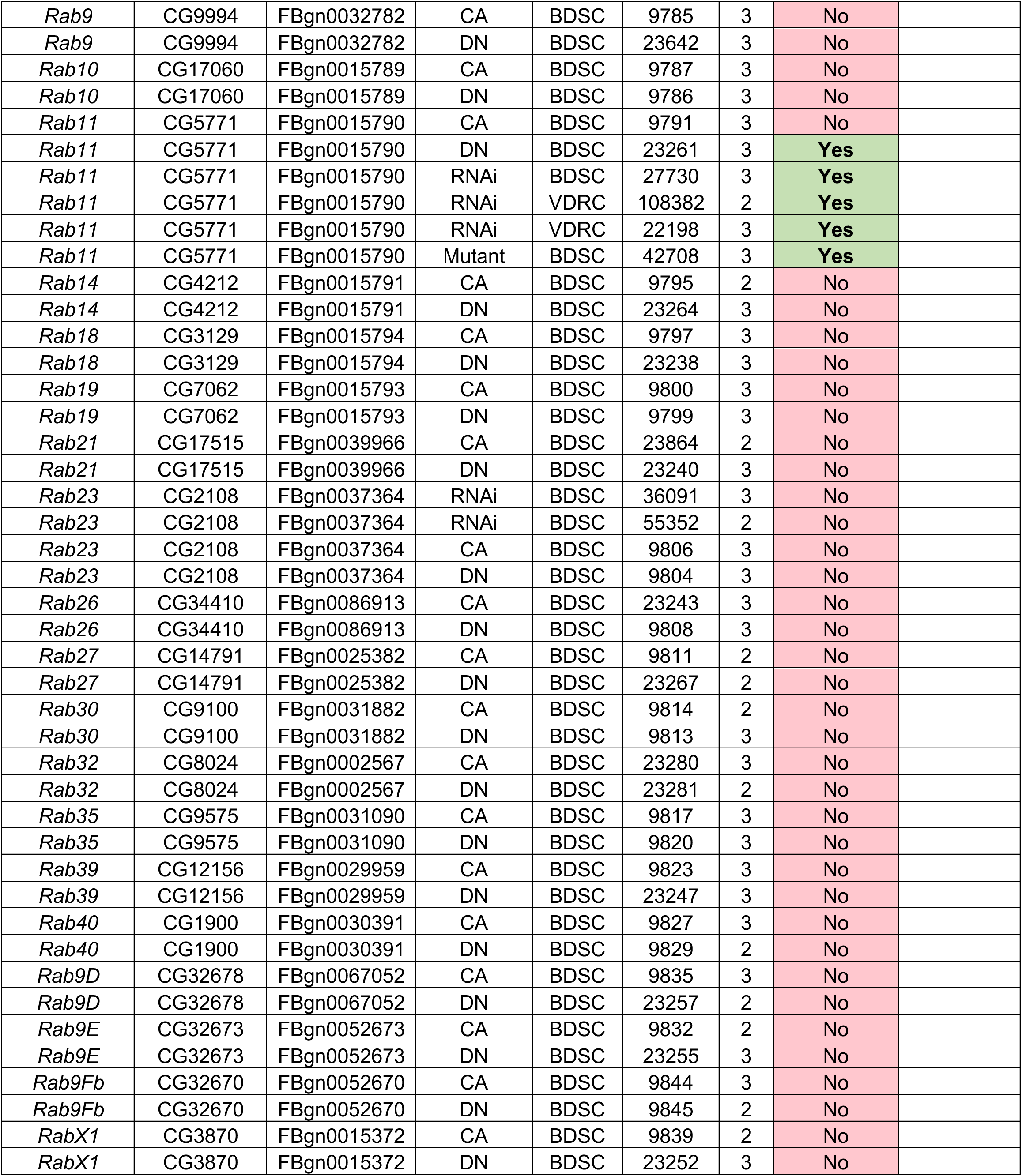
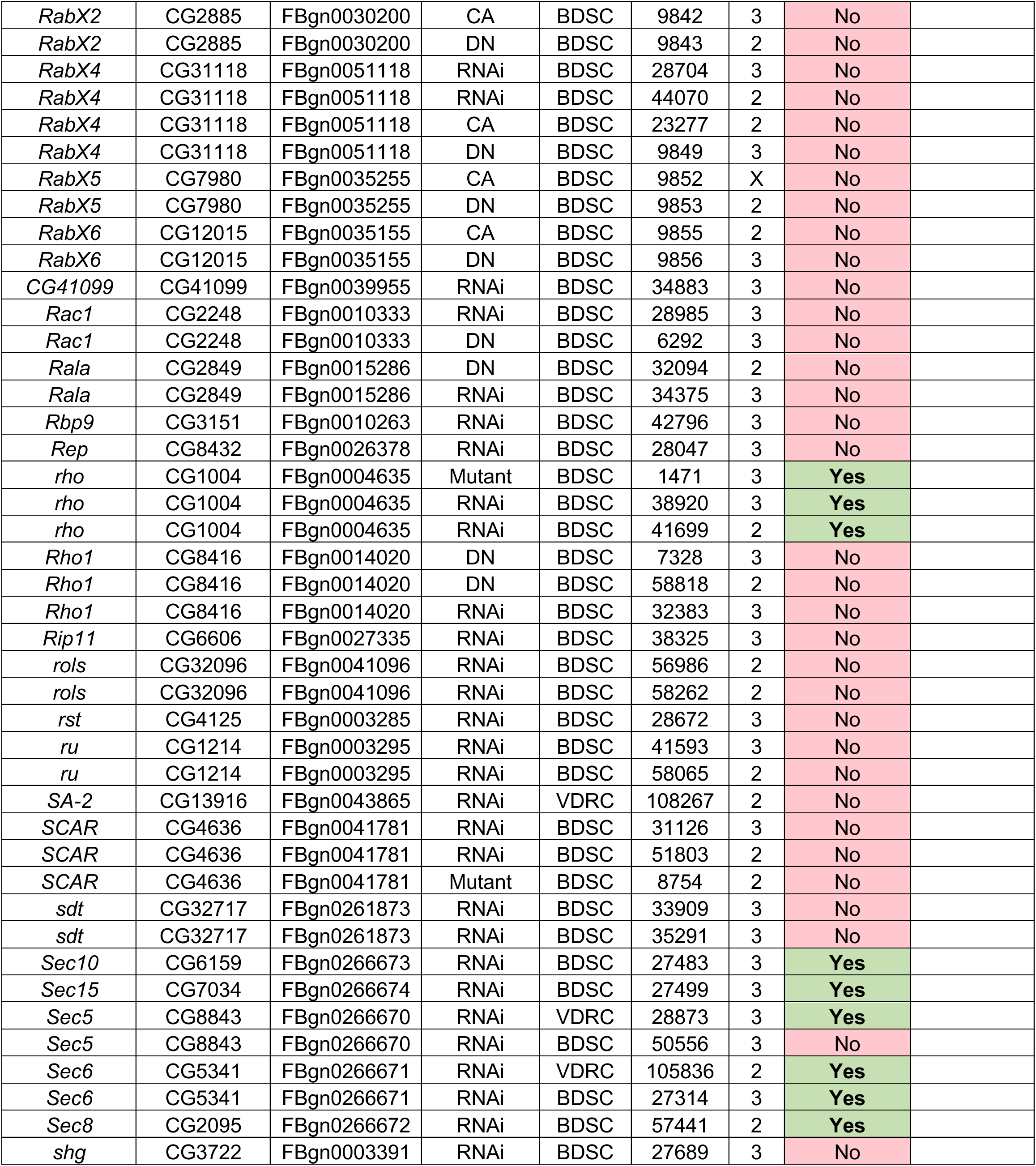
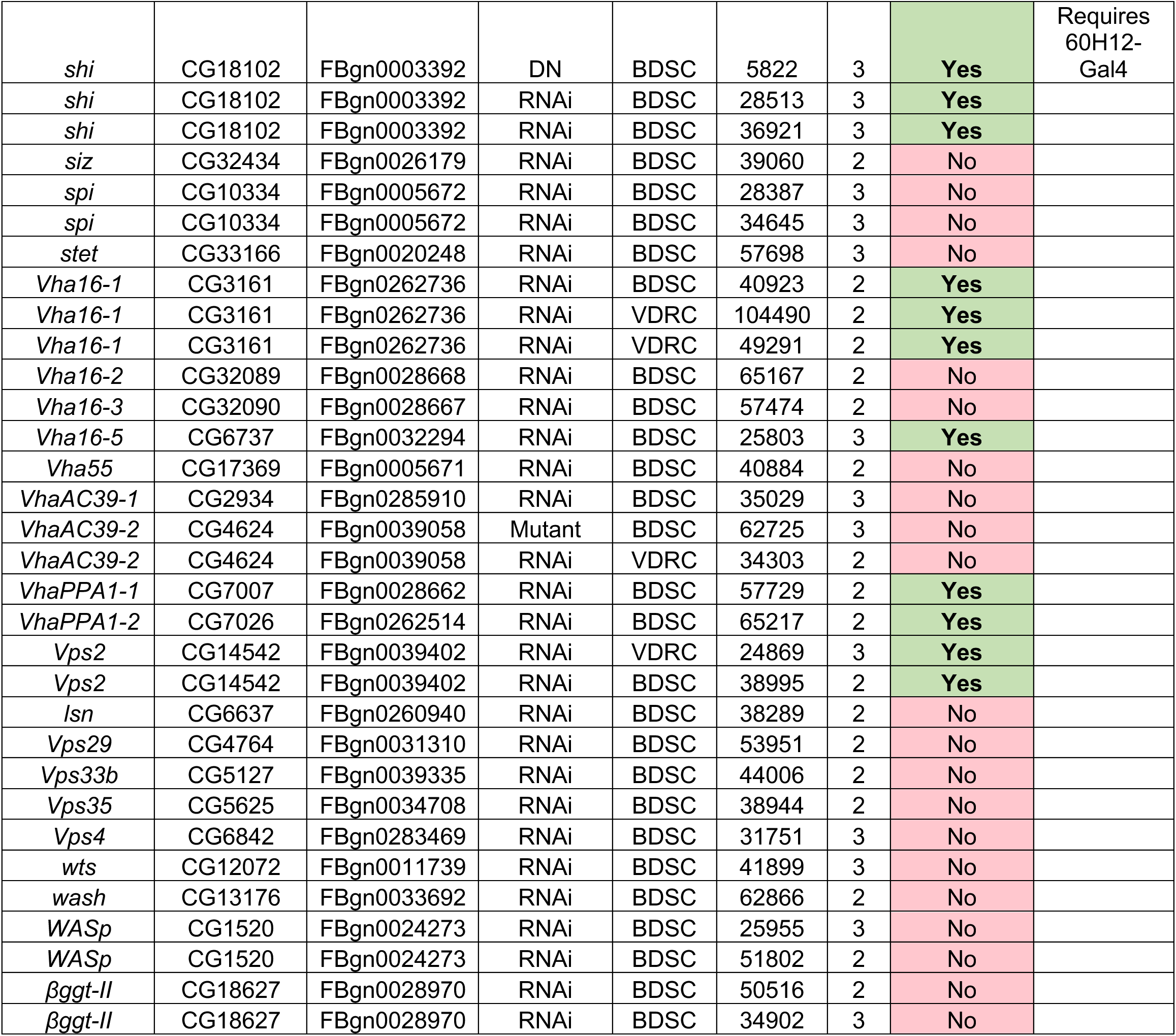
Primary and secondary candidate screen stock numbers used and results.

**Table S4.** Fly stocks used in addition to the screens.

| **Stock Name** | **Stock Number** | **Origin** | **References** |
| --- | --- | --- | --- |
| *w<sup>1118</sup>* | 3605 | BDSC |  |
| *tub-Gal4* | 5138 | BDSC |  |
| *tub-Gal80<sup>ts</sup>* | NA | NA |  |
| *UAS-dBrainbow* | 34513 | BDSC | (12) |
| *UAS-dBrainbow* | 34514 | BDSC | (12) |
| *Hsp70>cre* | 851 | BDSC |  |
| *UAS-fzr RNAi* | 25550 | VDRC | (14,15) |
| *UAS-shi RNAi #1* | 28513 | BDSC |  |
| *UAS-shi RNAi #2* | 36921 | BDSC |  |
| *UAS-Rab5 RNAi #1* | 30518 | BDSC |  |
| *UAS-Rab5 RNAi #2* | 67877 | BDSC |  |
| *UAS-Rab11 RNAi #1* | 27730 | BDSC |  |
| *UAS-Rab11 RNAi #2* | 22198 | VDRC |  |
| *UAS-SCAR RNAi #1* | 36121 | BDSC | (30) |
| *UAS-SCAR RNAi #2* | 51803 | BDSC | (31) |
| *UAS-kirre RNAi* | 27227 | VDRC | (32) |
| *UAS-sns RNAi* | 877 | VDRC | (32) |
| *UAS-schizo RNAi* | 36625 | VDRC | (33) |
| *UAS-sing RNAi* | 12202 | VDRC | (34) |
| *UAS-Cdc42<sup>DN</sup>* | 6288 | BDSC |  |
| *UAS-Rac1<sup>DN</sup>* | 6292 | BDSC |  |
| *UAS-Rho1<sup>DN</sup>* | 7328 | BDSC |  |
| *UAS-GFP-NLS* | 4776 | BDSC |  |
| *UAS-GFP-Myc-2x-FYVE* | 42712 | BDSC | (35, 36) |
| *UAS-YFP-Rab5* | 9775 | BDSC |  |
| *60H12-Gal4* | 39268 | BDSC |  |
| *UAS-shi<sup>DN</sup>* | 5811 | BDSC |  |
| *NrxIV-GFP* | 50798 | BDSC |  |
| *Df(1)BSC867* | 29990 | BDSC |  |
| *UAS-ogre RNAi* | 7136 | VDRC | (37, 38) |
| *byn-Gal4* | - | NA | (39) |
| *UAS-PA-GFP* | - | Lynn Cooley | (40) |
| *UAS-N<sup>DN</sup>* | - | NA | (41) |
| *UAS-shi-Venus* | - | Stefano De Renzis | (42) |
| *UAS-GFP-ogre* | - | Andrea Brand | (38) |

**Table S5.** Additional Methods.

| Panel | Additional Methods |
| --- | --- |
| 1D-D'' | *Hsp70>cre ; UAS-dBrainbow ; byn-Gal4* papillae dissected at 62 (D), 69 (D'), or 80 (D'') hours post-puparium formation (HPPF) at 25C. Hindguts were stained with Rabbit anti-GFP (Thermo-Fisher, A11122, 1:1000), Rat anti-HA (Sigma, 3F10, 1:100), and DAPI at 5µg/ml. |
| 1E | *Hsp70>cre ; UAS-dBrainbow ; byn-Gal4* papillae dissected at various HPPF at 25C. The area labelled by mKO2 was divided by total papillar area. |
| 1F | *Hsp70>cre ; UAS-dBrainbow ; byn-Gal4* papillae live-imaged at 69HPPF at 25C. |
| 1F' | Fluorescence intensity measured in neighboring cells during sharing onset (1F). |
| 1G-H | *byn-Gal4 / UAS-PA-GFP*, live-imaged during adulthood. Single secondary and primary cells were photoactivated and imaged every 3s. |
| 2A | UAS-RNAis and dominant-negative versions of 77 genes representing a wide range of cellular roles were screened (*Hsp70>cre ; UAS-dBrainbow ; byn-Gal4*) for sharing defects. Animals expressing both *UAS-dBrainbow* and an *UAS*-driven RNAi or mutant gene were raised at 25C and shifted to 29C at L3. If a given RNAi or DN line was lethal when expressed with the *byn-Gal4* driver, a *Gal80<sup>ts</sup>* was crossed in and the animals raised at 18C with a shift to 29C at pupation. Given the robustness of cytoplasmic sharing in WT animals, gene knockdowns or mutants with even single cell defects in sharing were considered "hits". |
| 2B | Secondary screen of 36 genes representing various categories of membrane trafficking (*Hsp70>cre ; UAS-dBrainbow ; byn-Gal4*) for sharing defects. Animals expressing both *UAS-dBrainbow* and an *UAS*-driven RNAi were raised at 25C and shifted to 29C at L3. If a given RNAi line was lethal when expressed with the *byn-Gal4* driver, a *Gal80<sup>ts</sup>* was crossed in and the animals raised at 18C with a shift to 29C at pupation. Given the robustness of cytoplasmic sharing in WT animals, gene knockdowns with even single cell defects in sharing were considered "hits". |
| 2C | Secondary screen (*Hsp70>cre ; UAS-dBrainbow ; byn-Gal4*) of dominant-negative and constitutively-active variants of the *Drosophila* Rab GTPases. *UAS-Rab11<sup>DN</sup>* and *UAS-Rab14<sup>DN</sup>* required a *Gal80<sup>ts</sup>* repressor and temperature shifts from 18C to 29C at pupation. *UAS-Rab1<sup>DN</sup>* and *UAS-Rab5<sup>DN</sup>* required papillar-specific expression using an alternative *Gal4* driver (*60H12-Gal4*), *Gal80<sup>ts</sup>* repressor, and temperature shifts from 18C to 29C at pupation. |
| 2D | *Hsp70>cre ; UAS-dBrainbow ; byn-Gal4, Gal80<sup>ts</sup>* animals dissected pre-sharing (48 HPPF at 29C). |
| 2D' | *Hsp70>cre ; UAS-dBrainbow ; byn-Gal4, Gal80<sup>ts</sup>* animals raised at 18C and shifted to 29C at pupation and dissected post-sharing (young adult). |
| 2E | Young adult animals expressing *UAS-shi RNAi #1* in a *Hsp70>cre ; UAS-dBrainbow ; byn-Gal4, Gal80<sup>ts</sup>* background. Animals were shifted from 18C to 29C at pupation to maximize RNAi and minimize animal lethality. |
| 2F | Young adult animals expressing *UAS-Rab5 RNAi #1* in a *Hsp70>cre ; UAS-dBrainbow ; byn-Gal4, Gal80<sup>ts</sup>* background. Animals were shifted from 18C to 29C at 1-2 days PPF to maximize RNAi and minimize animal lethality. |
| 2G | Young adult animals expressing *UAS-Rab11 RNAi #2* in a *Hsp70>cre* ; *UAS-dBrainbow* ; *byn-Gal4*, *Gal80<sup>ts</sup>* background. Animals were shifted from 18C to 29C at 1-2 days PPF to maximize RNAi and minimize animal lethality. |
| 2H | Animals were shifted and dissected as in 2D-G. Additionally, *Hsp70>cre* ; *UAS-dBrainbow* ; *byn-Gal4*, *Gal80<sup>ts</sup>* animals expressing *UAS-shi RNAi #2* were raised at 18C and shifted to 29C at pupation, animals expressing *UAS-Rab5 RNAi #2* were raised at 18C and shifted to 29C at L3, and animals expressing *UAS-Rab11 RNAi #1* were raised at 18C and shifted to 29C at 1-2 days PPF. |
| 3A-A' | Pupae expressing the early and late endosome marker *UAS-GFP-myc-2x-FYVE* were dissected pre (A, 48HPPF at 29C) and post (A', 72HPPF at 29C) sharing onset. |
| 3B | Pupae expressing *UAS-GFP-myc-2x-FYVE* in a *UAS-shi RNAi #1* background at a post sharing time point (24HPPF at 18C + 72 hours at 29C). |
| 3C | Aggregated line profiles of *UAS-GFP-myc-2x-FYVE* intensity across papilla. |
| 3D-D' | Pupae expressing *UAS-shi-Venus* were dissected pre (D, 48HPPF at 29C) and post (D', 72HPPF at 29C) sharing onset. |
| 3E | Aggregated line profiles of Shi-Venus intensity from the basal (0% distance) to the apical (100% distance) edges of the papilla. See 3C. |
| 3F-F'' | Transmission electron micrographs of the microvillar-like structures of pupal papillae pre (F, 60HPPF at 25C), mid (F', 66HPPF at 25C), and post (F'', 69HPPF at 25C) cytoplasm sharing onset. |
| 3G-G'' | Electron micrographs of mitochondria and surrounding membrane material pre (G, 60HPPF at 25C), mid (G', 66HPPF at 25C), and post (G'', 69HPPF at 25C) |
| 3H | Electron micrograph of microvillar-like structures of WT (*w<sup>1118</sup>*) young adult papillar cells. |
| 3I | Electron micrograph of microvillar-like structures of young adult *byn-Gal4*, *Gal80<sup>ts</sup>* > *UAS-shi RNAi #2* (raised at 18C, shifted at pupation to 29C). |
| 3J | Electron micrograph of microvillar-like structures of young adult *byn-Gal4*, *Gal80<sup>ts</sup>* > *UAS-Rab5 RNAi #1* animals (raised at 18C, shifted at 1-2 days PPF to 29C). |
| 3K | Electron micrograph of mitochondria and surrounding membrane material of WT (*w<sup>1118</sup>*) young adult papillar cells. |
| 3L | Electron micrograph of mitochondria and surrounding membrane material of young adult *byn-Gal4*, *Gal80<sup>ts</sup>* > *UAS-shi RNAi #2* (raised at 18C, shifted at pupation to 29C). |
| 3M | Electron micrograph of mitochondria and surrounding membrane material of young adult *byn-Gal4*, *Gal80<sup>ts</sup>* > *UAS-Rab5 RNAi #1* animals (raised at 18C, shifted at 1-2 days PPF to 29C). |
| 3N | Electron micrograph of post-sharing WT (TM3 / *UAS-shi RNAi #1*) pupa (24HPPF at 18C, shifted to 29C for 50 hours, then dissected) |
| 3O | Electron micrograph of post-sharing *byn-Gal4*, *Gal80<sup>ts</sup>* > *UAS-shi RNAi #1* pupa (24HPPF at 18C, shifted to 29C for 50 hours, then dissected) |
| 3P | Gap junction length / (gap junction length + septate junction length) measured in WT and *UAS-shi RNAi #1* pupae (see 3N-3O). Each point represents an image of a junction. |
| 4A-A'' | Electron micrographs of apical junctions (adherens, septate, and gap) pre (A, 60HPPF at 25C), mid (A', 66HPPF at 25C), and post (A'', 69HPPF at 25C) |
| 4B | Gap junction length / (gap junction length + septate junction length) measured in pupae pre (60HPPF at 25C), mid (66HPPF at 25C), and post (69HPPF at 25C) sharing onset. Each point represents an image of a junction. |
| 4C | Relative innexin transcript abundance (innexin X transcripts / total innexin transcripts) using data from Fly Atlas 2 (25) and RNA-seq of adult *w<sup>1118</sup>* rectums performed in the Fox Lab. |
| 4D-D' | Pupae with endogenously GFP-tagged *NrxIV* (*NrxIV-GFP*) dissected pre (D, 48HPPF) and post (D', 72HPPF) sharing onset. |
| 4E-E' | Pupae stained with *Inx3* antibody (gift from Reinhard Bauer, rabbit, 1:75) pre (D, 48HPPF) and post (D', 58HPPF, papillae do not stain well at later timepoints) sharing onset. |
| 4F | Young adult animals expressing no transgene (WT), *UAS-ogre<sup>DN</sup>*, *UAS-ogre RNAi*, or containing a deficiency covering *ogre*, *Inx2*, and *Inx7* in a *Hsp70>cre* ; *UAS-dBrainbow* ; *byn-Gal4*, *Gal80<sup>ts</sup>* background. Animals were raised at 25C until L3 and then shifted to 29C until dissection at young adulthood. |
| 4G | See 4F. |
| 4H | *60H12-Gal4*, *Gal80<sup>ts</sup>* driving *UAS-shi<sup>DN</sup>* and WT siblings were shifted from 18C to 29C at pupation. *byn-Gal4*, *Gal80<sup>ts</sup>* driving *UAS-ogre<sup>DN</sup>* animals and WT siblings were raised at 25C and shifted to 29C at L3. Animals 1-3 days post-eclosion were sorted into sex-matched groups and fed a control diet or a high salt (2% NaCl) diet. Survival was assessed once per day for 10 days. |
| S1A | *Hsp70>cre* ; *UAS-dBrainbow* ; *tubulin-Gal4* animals raised at 29C. Tissues dissected at adulthood. |
| S1E | *Hsp70>cre* ; *UAS-dBrainbow* ; *byn-Gal4* animals were shifted from 25C to 29C during L3 and dissected at adulthood. |
| S1F | *Hsp70>cre* ; *UAS-dBrainbow* / *UAS-fzr RNAi* ; *byn-Gal4* animals were shifted from 25C to 29C during L2 to maximize *fzr* knock down during endocycling. Animals were dissected at adulthood. |
| S1G | *Hsp70>cre* ; *UAS-dBrainbow* ; *byn-Gal4* / *UAS-N<sup>DN</sup>* animals were shifted from 25C to 29C during L3 to ensure maximum *UAS-N<sup>DN</sup>* expression during mitoses. Animals were dissected at adulthood. |
| S2A | *Hsp70>cre* ; *UAS-dBrainbow* ; *byn-Gal4*, *Gal80<sup>ts</sup>* animals expressing various previously published myoblast fusion RNAis raised at 25C and shifted to 29C at L3 and dissected post-sharing (young adult). |
| S2B | *Hsp70>cre* ; *UAS-dBrainbow* ; *byn-Gal4*, *Gal80<sup>ts</sup>* animals expressing various previously published UAS-dominant negative active regulators raised at 18C and shifted to 29C at L3 and dissected post-sharing (young adult). |
| S2C | Papillar cells were identified using *byn-Gal4*, *Gal80<sup>ts</sup>*, driving *UAS-GFP<sup>NLS</sup>* expression. Cells were counted in one, z-sectioned half of the papillae and multiplied by 2 to give an approximate cell count. |
| S2D | *Hsp70>cre* ; *UAS-dBrainbow* ; *byn-Gal4*, *Gal80<sup>ts</sup>* animals were raised at 18C until 3-4 days PPF and shifted to 29C and dissected at young adulthood. |
| S2E | *Hsp70>cre* ; *UAS-dBrainbow* ; *byn-Gal4*, *Gal80<sup>ts</sup>* animals expressing *UAS-shi RNAi #1* were raised at 18C until 3-4 days PPF and shifted to 29C and dissected at young adulthood. |
| S3A | See 3A-3C. Basal and apical membrane defined as 10-20% and 90-100% total distance of papillae, respectively. |
| S3B-B' | *byn-Gal4 > UAS-Rab5-YFP* animals dissected pre (48HPPF, 29C) and post (72HPPF, 29C) sharing onset. |
| S3C | See S3B-B' and 3C. |
| S3D-D'' | Electron micrographs of apical junctions (adherens, septate, and gap) pre (D, 60HPPF at 25C), mid (D', 66HPPF at 25C), and post (D'', 69HPPF at 25C) |
| S3E | Electron micrograph of apical junctions (adherens, septate, and gap) of WT (*w<sup>1118</sup>*) young adult papillar cells. |
| S3F | Electron micrograph of apical junctions (adherens, septate, and gap) of young adult *byn-Gal4, Gal80<sup>ts</sup> > UAS-shi RNAi #2* (raised at 18C, shifted at pupation to 29C). |
| S3G | Electron micrograph of apical junctions (adherens, septate, and gap) of young adult *byn-Gal4, Gal80<sup>ts</sup> > UAS-Rab5 RNAi #1* animals (raised at 18C, shifted at 1-2 days PPF to 29C). |
| S3H | See 3N-O. Junction width was measured throughout and averaged per image. Each point represents one image of a junction. |
| S3H' | See 3N-O. Junction width was measured throughout and averaged per image. Each point represents one image of a junction. |
| S3H'' | See 3N-O. Raw lengths shown were used to calculate "fraction gap junction" in 3P. Each point represent one image of a junction. |
| S4A | TEM of young adult (*w<sup>1118</sup>*) papilla. |
| S5A | See 4A-B. Junction width was measured throughout and averaged per image. Each point represents one image of a junction. |
| S5A' | See 4A-B. Junction width was measured throughout and averaged per image. Each point represents one image of a junction. |
| S5A'' | See 4A-B. Raw lengths shown were used to calculate "fraction gap junction" in 3P. Each point represent one image of a junction. |
| S5B-B' | Pupae expressing *byn-Gal4, Gal80<sup>ts</sup> > UAS-ogre<sup>DN</sup> (UAS-GFP-ogre)* dissected pre (B, 48HPPF, 29C) and post (B', 72HPPF, 29C) sharing onset. |
| S5C | *byn-Gal4 > UAS-GFP<sup>NLS</sup>* dissected pre (48HPPF, 29C) sharing onset. |
| S5D | *60H12-Gal4 > UAS-GFP<sup>NLS</sup>* dissected pre (48HPPF, 29C) sharing onset. The pan-hindgut driver used in previous experiments, *brachyenteron (byn>Gal4)*, causes animal lethality with *shi*, *Rab5*, and *Rab11* knockdown within a few days. We therefore screened for and identified an alternative, papillae-specific driver (*60H12>Gal4*), derived from regulatory sequences of the hormone receptor gene *Proctolin Receptor*. *60H12>shi<sup>DN</sup>* animals are viable on a control diet allowing us to test papillar function on a high-salt diet. |
| S5E | *Hsp70>cre ; UAS-dBrainbow ; 60H12-Gal4* animals raised at 18C and shifted to 29C at pupation and dissected as young adults. |
| S5F | *Hsp70>cre ; UAS-dBrainbow ; 60H12-Gal4 / UAS-shi<sup>DN</sup>* animals raised at 18C and shifted to 29C at pupation and dissected as young adults. |
| S5G | See S5E-F. |

**Table S6.**
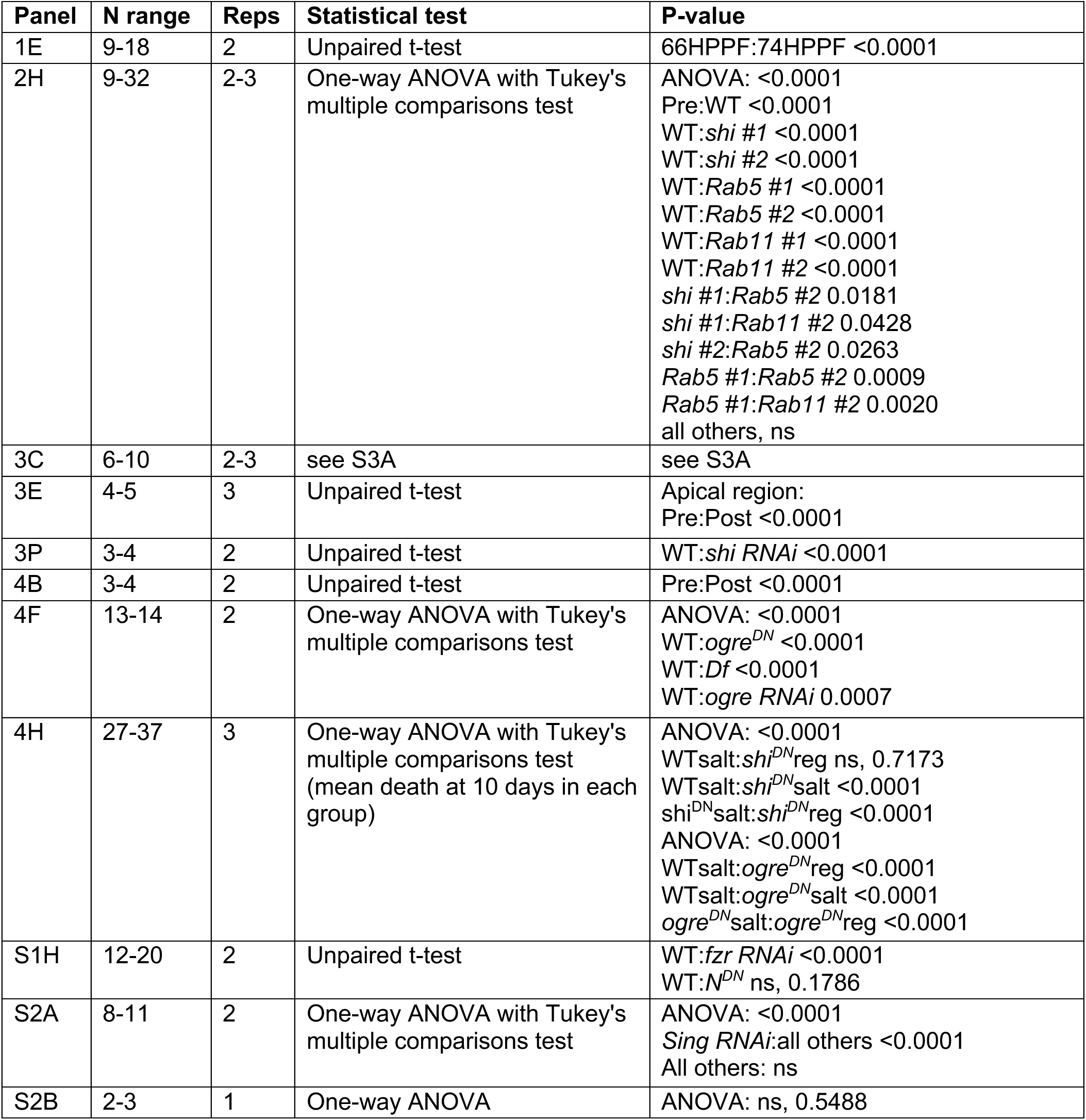
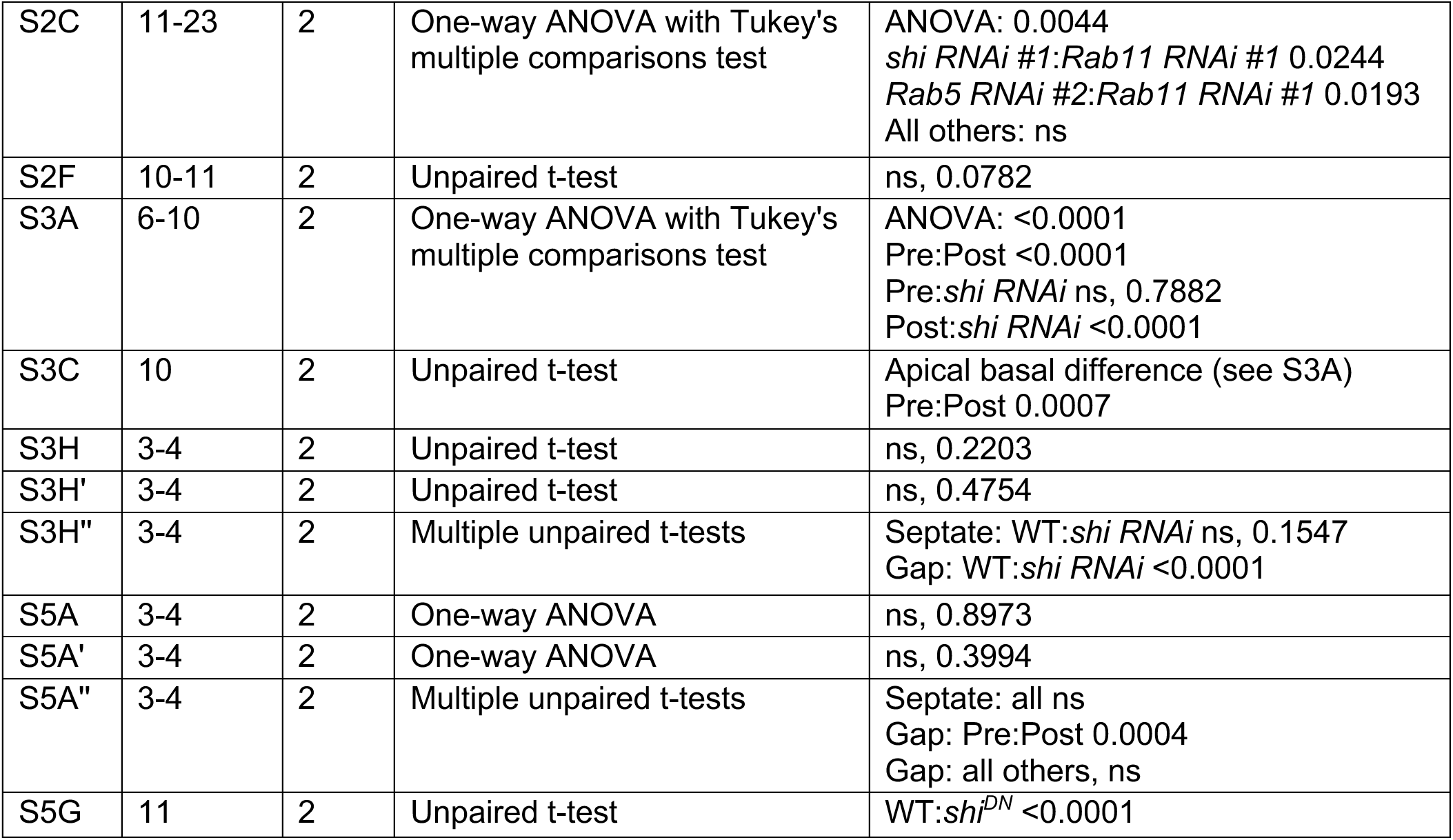
Additional statistics.

| Panel | N range | Reps | Statistical test | P-value |
| --- | --- | --- | --- | --- |
| 1E | 9-18 | 2 | Unpaired t-test | 66HPPF:74HPPF <0.0001 |
| 2H | 9-32 | 2-3 | One-way ANOVA with Tukey's multiple comparisons test | ANOVA: <0.0001<br>Pre:WT <0.0001<br>WT:*shi* #1 <0.0001<br>WT:*shi* #2 <0.0001<br>WT:*Rab5* #1 <0.0001<br>WT:*Rab5* #2 <0.0001<br>WT:*Rab11* #1 <0.0001<br>WT:*Rab11* #2 <0.0001<br>*shi* #1:*Rab5* #2 0.0181<br>*shi* #1:*Rab11* #2 0.0428<br>*shi* #2:*Rab5* #2 0.0263<br>*Rab5* #1:*Rab5* #2 0.0009<br>*Rab5* #1:*Rab11* #2 0.0020<br>all others, ns |
| 3C | 6-10 | 2-3 | see S3A | see S3A |
| 3E | 4-5 | 3 | Unpaired t-test | Apical region:<br>Pre:Post <0.0001 |
| 3P | 3-4 | 2 | Unpaired t-test | WT:*shi* RNAi <0.0001 |
| 4B | 3-4 | 2 | Unpaired t-test | Pre:Post <0.0001 |
| 4F | 13-14 | 2 | One-way ANOVA with Tukey's multiple comparisons test | ANOVA: <0.0001<br>WT:*ogre*<sup>DN</sup> <0.0001<br>WT:*Df* <0.0001<br>WT:*ogre* RNAi 0.0007 |
| 4H | 27-37 | 3 | One-way ANOVA with Tukey's multiple comparisons test (mean death at 10 days in each group) | ANOVA: <0.0001<br>WTsalt:*shi*<sup>DN</sup>reg ns, 0.7173<br>WTsalt:*shi*<sup>DN</sup>salt <0.0001<br>*shi*<sup>DN</sup>salt:*shi*<sup>DN</sup>reg <0.0001<br>ANOVA: <0.0001<br>WTsalt:*ogre*<sup>DN</sup>reg <0.0001<br>WTsalt:*ogre*<sup>DN</sup>salt <0.0001<br>*ogre*<sup>DN</sup>salt:*ogre*<sup>DN</sup>reg <0.0001 |
| S1H | 12-20 | 2 | Unpaired t-test | WT:*fzr* RNAi <0.0001<br>WT:*N*<sup>DN</sup> ns, 0.1786 |
| S2A | 8-11 | 2 | One-way ANOVA with Tukey's multiple comparisons test | ANOVA: <0.0001<br>*Sing* RNAi:all others <0.0001<br>All others: ns |
| S2B | 2-3 | 1 | One-way ANOVA | ANOVA: ns, 0.5488 |
| S2C | 11-23 | 2 | One-way ANOVA with Tukey's multiple comparisons test | ANOVA: 0.0044<br>*shi RNAi* #1:*Rab11 RNAi* #1 0.0244<br>*Rab5 RNAi* #2:*Rab11 RNAi* #1 0.0193<br>All others: ns |
| S2F | 10-11 | 2 | Unpaired t-test | ns, 0.0782 |
| S3A | 6-10 | 2 | One-way ANOVA with Tukey's multiple comparisons test | ANOVA: <0.0001<br>Pre:Post <0.0001<br>Pre:*shi RNAi* ns, 0.7882<br>Post:*shi RNAi* <0.0001 |
| S3C | 10 | 2 | Unpaired t-test | Apical basal difference (see S3A)<br>Pre:Post 0.0007 |
| S3H | 3-4 | 2 | Unpaired t-test | ns, 0.2203 |
| S3H' | 3-4 | 2 | Unpaired t-test | ns, 0.4754 |
| S3H'' | 3-4 | 2 | Multiple unpaired t-tests | Septate: WT:*shi RNAi* ns, 0.1547<br>Gap: WT:*shi RNAi* <0.0001 |
| S5A | 3-4 | 2 | One-way ANOVA | ns, 0.8973 |
| S5A' | 3-4 | 2 | One-way ANOVA | ns, 0.3994 |
| S5A'' | 3-4 | 2 | Multiple unpaired t-tests | Septate: all ns<br>Gap: Pre:Post 0.0004<br>Gap: all others, ns |
| S5G | 11 | 2 | Unpaired t-test | WT:*shi<sup>DN</sup>* <0.0001 |

## SUPPLEMENTAL FIGURE LEGENDS

**Figure S1.**
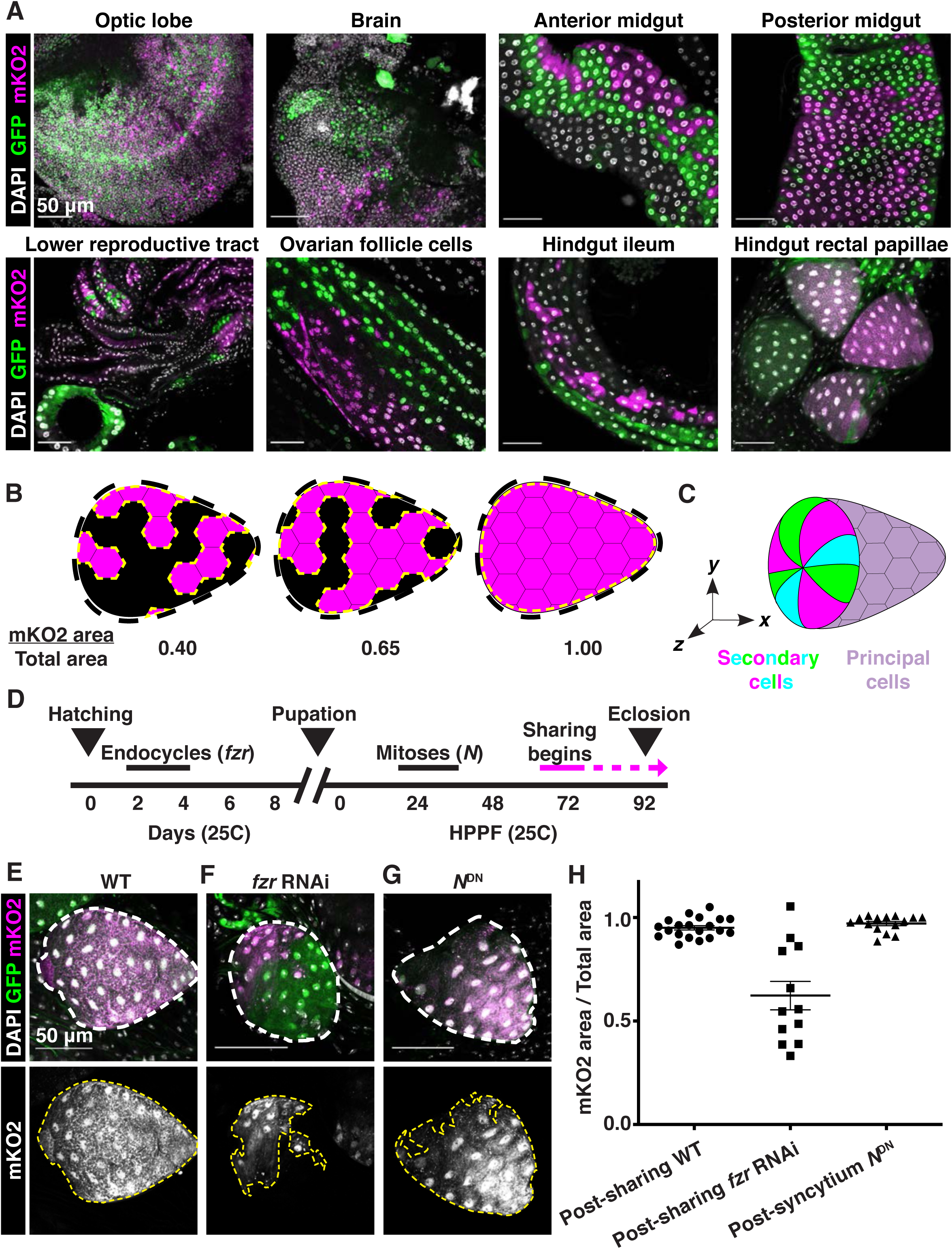
The hindgut rectal papillae share cytoplasm independent of mitosis. (**A**) Representative images of dBrainbow expression in the indicated adult tissues. (**B**) Schematic of cytoplasmic sharing quantification. The mKO2-positive papillar area is divided by the total papillar area to give a score of cytoplasmic sharing. Numbers close to 1 indicate near-complete sharing. (**C**) Schematic of principal cells (sharing) and secondary cells (non-sharing) at the papillar base that together form each papilla. (**D**) Approximate timeline of cytoplasm sharing onset (68-74 HPPF) within papillar development (*14*). Cytoplasmic sharing is temporally separate from papillar mitoses. (**E-G**) Representative adults expressing dBrainbow in a (**E**) wild-type (WT), (**F**) *fzr RNAi* (p<0.0001), or (**G**) *N^DN^* background (p=0.8786). (**H**) Quantification of cytoplasmic sharing in adult WT, *fzr RNAi*, and *N^DN^*-expressing animals (N=12-20, rep=2).

**Figure S2.**
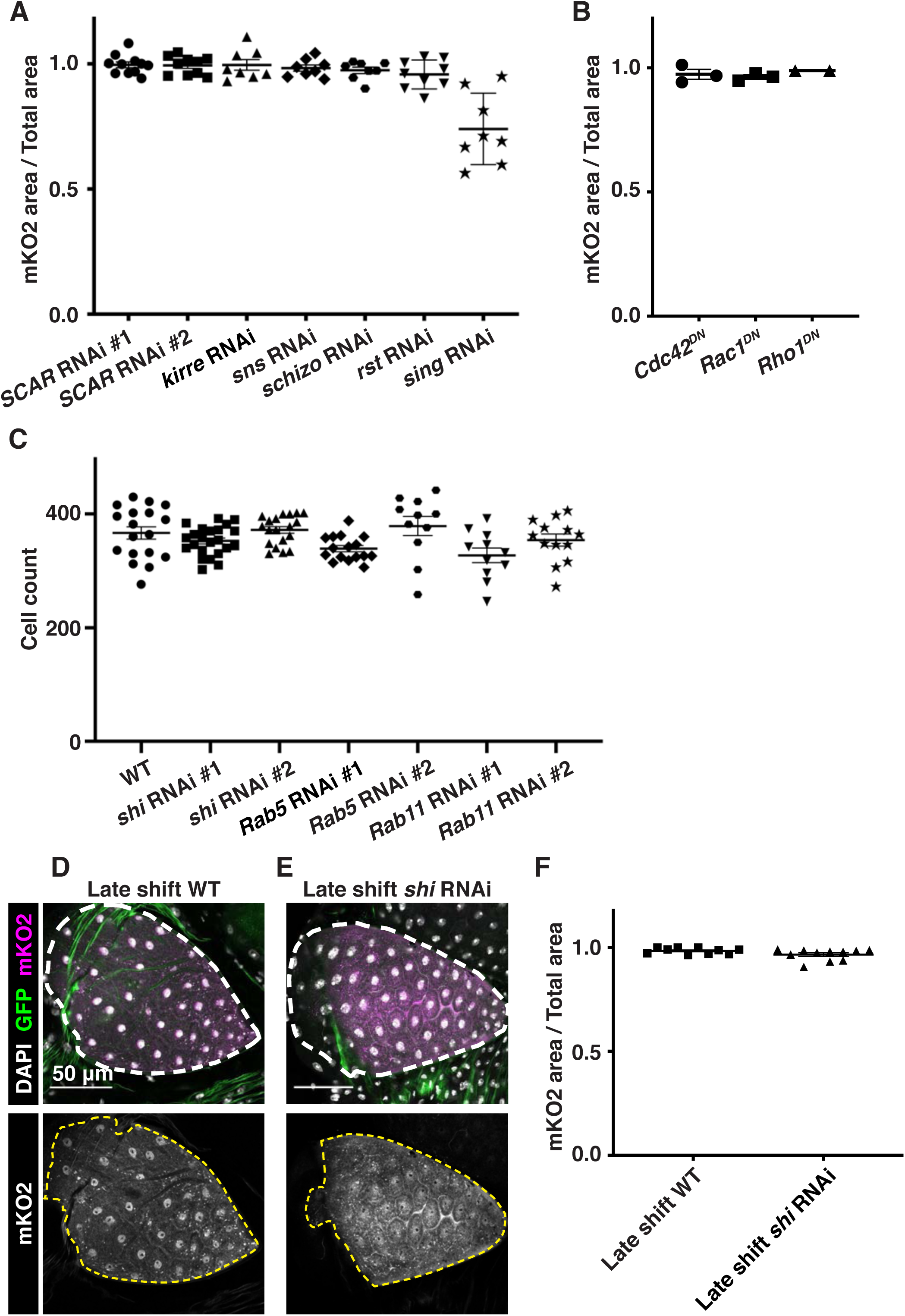
Membrane trafficking genes expressed during a developmental window regulate cytoplasm sharing. (**A**) Quantification of cytoplasmic sharing in animals expressing dsRNA for myoblast fusion regulators (N=8-11, rep=2). All knockdown lines are previously published (*30–34*). Only *sing RNAi* significantly differs from WT (p<0.0001). (**B**) Quantification of cytoplasmic sharing in animals expressing dsRNA for Rho family GTPases. (**C**) Cell counts in WT and knockdown rectal papillae (N=11-23, rep=2). Only *Rab11 #1 RNAi* had a significantly different cell number than WT (p=0.0323). (**D-E**) Representative animals expressing *dBrainbow* in either a WT (**D**) or *shi RNAi* (**E**) genetic background were raised at 18C until 3-4 days PPF and shifted to 29C to induce *shi* knockdown at a later timepoint than in Figure 2E**, 2H**. (**F**) Sharing quantification in late-induced animals (N=10-11, rep=2).

**Figure S3.**
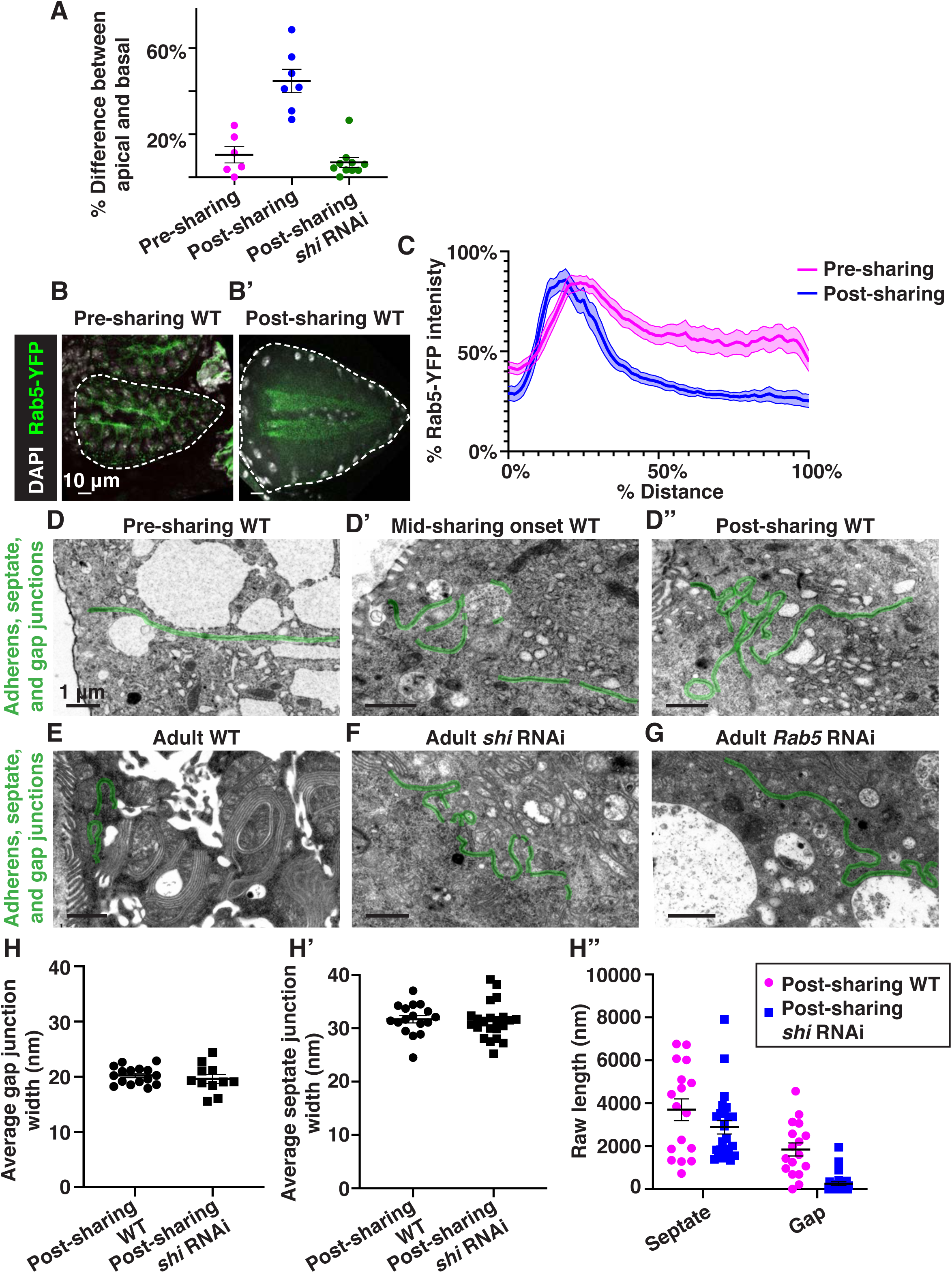
Changes in endosome polarity and apical junction shape accompany the onset of cytoplasm sharing. (**A**) Quantification of the average endosome intensity difference between representative basal and apical areas across papillae in Figure 3A-C (N=6-10, rep=2). (**B-B’**) Representative localization of Rab5-YFP, green, before sharing onset (**B**) and after sharing onset (**B’**). (**C**) Aggregated line profiles of Rab5-YFP intensity before and after the beginning of sharing (N=10, rep=2). (**D-D’’**) Representative TEMs of apical (adherens, septate, and gap) junctions pre (**D**), mid (**D’**), and post (**D’’**) sharing onset. (**E-G**) Representative TEMs of apical junctions of post-sharing adult WT (**E**), *shi RNAi* (**F**), and *Rab5 RNAi* (**G**) papillar cells. (**H-H’’**) Apical junction electron micrograph measurements of post-sharing WT and *shi RNAi* pupal papillar cells (N=3-4, rep=2). Average gap junction (**H**) and septate junction (**H’**) widths were measured alongside gap and septate junction length. Width measurements were taken along the length of each cell-cell junction and averaged to give one point per cell-cell junction. (**H’’**) Raw septate and gap junction lengths (nm) that were used to calculate gap junction ratio in Figure 3P.

**Figure S4.**
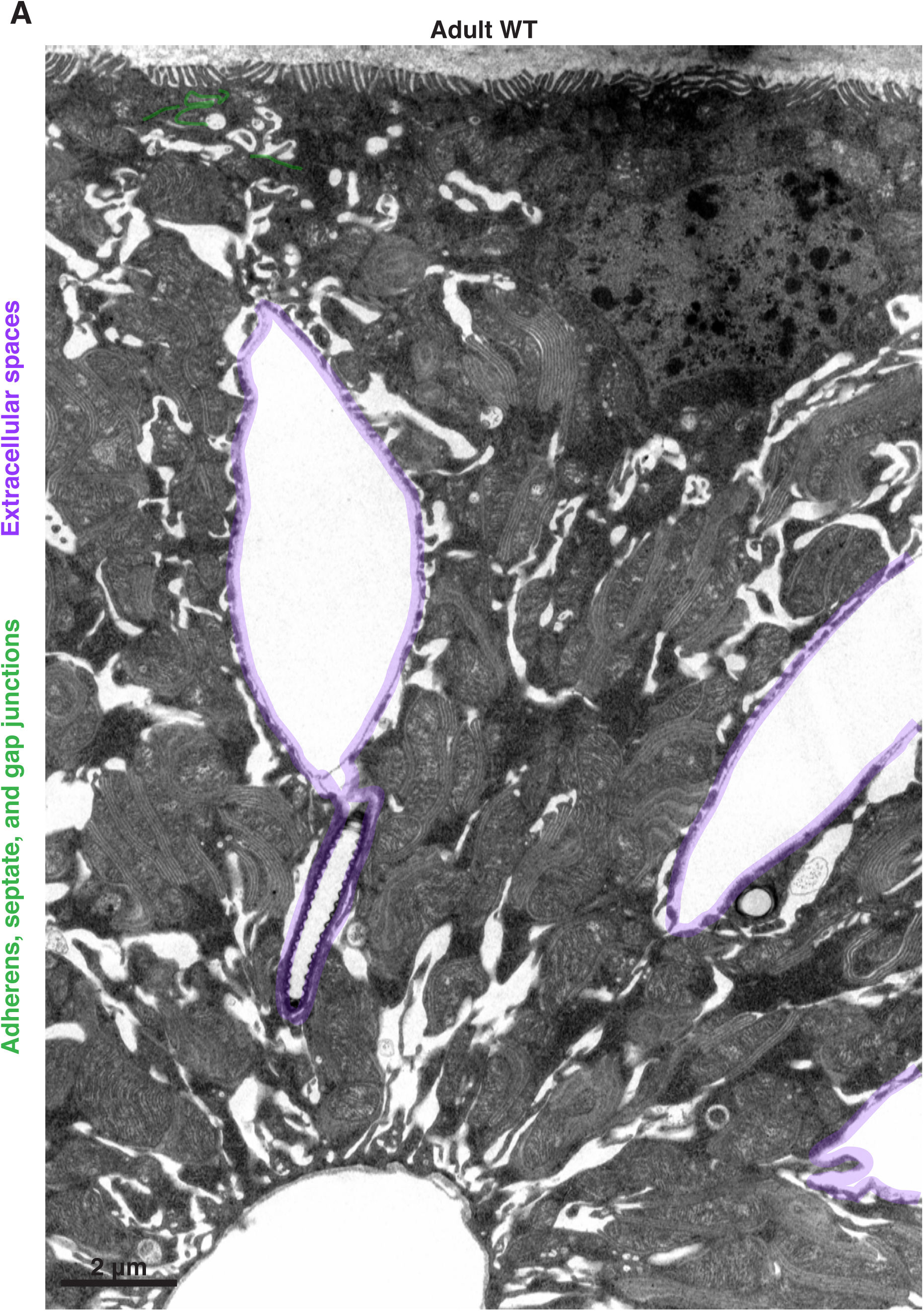
Extracellular spaces separate nuclei throughout much of the papillar lateral membrane. (**A**) Representative TEM cross-section of an adult WT papilla.

**Figure S5.**
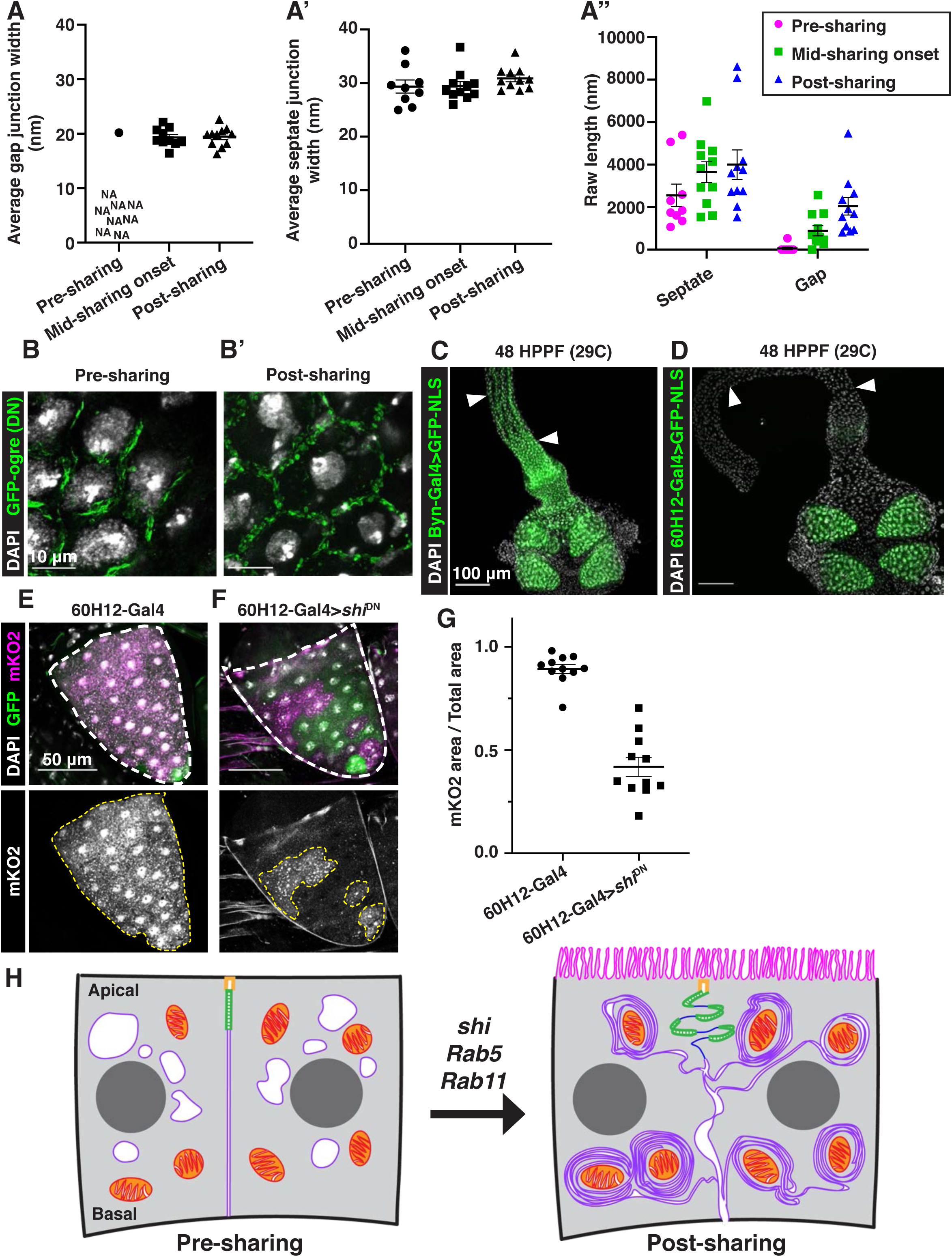
Gap junction formation coincides with cytoplasm sharing onset. (**A-A’’**) Apical junction TEM measurements of pre, mid, and post-sharing onset pupal papillar cells (N=3-4, rep=2). Average gap junction (**A**) and septate junction (**A’**) widths were measured alongside gap and septate junction length. (**A’’**) Raw septate and gap junction lengths (nm) used to calculate gap junction ratio in Figure 4B. (**B-B’**) Gap junction localization visualized by *UAS-GFP-ogre* in pre (**B**) and post (**B’**) sharing onset pupae. (**C**) Representative image of *byn-Gal4* driving *GFP^NLS^* expression throughout the pre-sharing hindgut. (**D**) *60H12-Gal4* driving *GFP^NLS^* expression in pre-sharing papillae but not in the ileum or pylorus. (**E**) Representative image of *60H12-Gal4* driving dBrainbow in adult papillae. (**F**) Representative image of *60H12-Gal4* driving *shi^DN^* expression in a dBrainbow background in adult papillae. (**G**) Quantification of cytoplasm sharing in *60H12-Gal4* and *60H12-Gal4>shi^DN^* animals (N=11, rep=2). (**H**) Model of membrane and junctional changes requiring membrane trafficking genes that coincide with the onset of cytoplasm sharing.

